# Complementary frontoparietal and corticothalamic contributions to relational reasoning

**DOI:** 10.64898/2026.07.03.736406

**Authors:** Conor N. Robinson, Luke J. Hearne, Kartik K. Iyer, Takuya Ito, James A. Roberts, Luca Cocchi

**Affiliations:** Max Planck Institute for Human Cognitive and Brain Sciences, Leipzig, Germany; Clinical Brain Networks Group, QIMR Berghofer, Brisbane, QLD, Australia; Faculty of Health, Medicine and Behavioural Sciences, University of Queensland, Brisbane, QLD, Australia; Brain Modelling Group, QIMR Berghofer, Brisbane, QLD, Australia; T.J. Watson Research Center, IBM Research, Yorktown Heights, NY, USA

**Keywords:** frontoparietal, thalamus, reasoning, neural field, aperiodic, corticothalamic, complexity, cognitive demand

## Abstract

Complex reasoning requires frontal, parietal, and thalamic systems to manage increasing relational demands, yet the underlying circuit mechanisms remain unclear. Here, we combined EEG with biologically grounded corticothalamic neural field modelling while participants solved relational problems of graded complexity. Successful reasoning was associated with dissociable frontoparietal dynamics. Frontal regions showed increased theta-band power, whereas parietal regions showed reduced alpha- and beta-band power. Neural field modelling linked these dynamics to complementary complexity-dependent circuit adaptations. Parietal regions showed modulation of intracortical and corticothalamic gains, intrathalamic inhibition, prolonged loop delays, and faster synaptic filtering, whereas frontal regions primarily adjusted intracortical gains in a manner consistent with maintaining local excitatory–inhibitory balance and supporting longer temporal integration windows. Together, these empirical and model-derived insights demonstrate that problem-solving under increasing reasoning demand relies not simply on greater frontoparietal engagement, but on additional, region-specific reconfigurations of cortical and corticothalamic circuits.

## INTRODUCTION

The frontoparietal network (FPN) serves as a central hub for integrating information required for complex problem-solving and reasoning ^1,2^. Across a wide range of cognitively demanding tasks, increasing task difficulty is accompanied by heightened activity and reconfiguration of functional connectivity within frontal and parietal cortices ^3–7^. While these findings establish the central role of the FPN in reasoning, we still have limited insight into the temporal, spectral, and circuit-level mechanisms by which relations are integrated, particularly as relational demands increase.

Relational complexity (RC) provides a principled framework for quantifying reasoning demands, defined as the maximum number of relations that must be integrated within a single cognitive operation ^8^. Across diverse reasoning domains, including non-verbal visuospatial puzzles, propositional logic, transitive inference, and sentence comprehension tasks, RC functions as a domain-general marker of processing capacity limits ^8–12^. Accordingly, increases in relational complexity reliably impair performance and recruit frontoparietal regions and networks, linking graded behavioural costs to the neural engagement required for relational integration ^4,5^.

The Latin Square Task (LST) is a validated paradigm for manipulating RC ^11,13,14^. In the LST, participants solve single-constraint visuospatial puzzles in which relational demands are systematically varied. A prior fMRI–EEG study using the LST identified a 2–4.2 s post-stimulus window in which the FPN supports relational-complexity integration ^15^. These delayed representations are distinct from early stimulus-locked responses and aligned with high-order computational strategies learned by artificial neural networks trained on the same task ^16,17^. Oscillatory activity in the theta, alpha, and beta bands has been linked to both local processing and long-range coordination across frontoparietal systems during higher-order cognition ^18,19^ and recent work has begun to characterise the role of frequency-specific neural dynamics in supporting reasoning under increasing cognitive demands ^20–22^. However, the mechanisms coordinating integration within this temporal window remain unresolved.

Corticothalamic interactions provide a plausible mechanism through which modulation of FPN activity can be achieved to facilitate the processing of RC ^23–25^. The thalamus is anatomically well-positioned to support the modulation of cortical integration, given its organisation and extensive cortical connectivity ^23,24^. While the thalamus is involved during relational processing ^26,27^, progressive increases in RC do not reliably produce corresponding increases in mean thalamic activation ^1^. This functional dissociation suggests that complexity-related effects may be mediated by distinct changes in thalamo-FPN coordination or gain, rather than by overall increases in thalamic activity. Increasing relational demands may thus be supported by thalamic-induced selective amplification of neural activity in frontal or parietal cortices, or by distinct reconfigurations of corticothalamic loops that shape frontoparietal dynamics. Neural field theory (NFT) models provide a mechanistic framework for addressing these questions by linking macroscopic EEG spectra to biologically grounded circuit parameters governing cortico–cortical, corticothalamic, and intrathalamic interactions ^28–35^.

Here, we used EEG and corticothalamic NFT to investigate how frontal, parietal, and thalamic circuits support relational integration as reasoning complexity increases. Focusing on the established LST RC integration window ^15^ we characterise complexity-dependent changes in local spectral power and frequency-specific frontoparietal synchronisation, and their relationships with behaviour. We then apply NFT modelling to dissociate cortical and corticothalamic contributions to these effects. Our findings reveal a functional dissociation between frontal and parietal processing as complexity increases, characterised by region-specific adjustments in the excitation–inhibition balance and corticothalamic gain. Together, these results advance mechanistic insight into how the frontoparietal–thalamic system accommodates increasingly complex relational processes supporting human reasoning.

## RESULTS

We started by assessing participants’ behavioural performance. We modified the puzzle format to accommodate shorter presentation and response times, while keeping stimuli within the participant’s visual field (Figure 1A). Each trial presented a 4×4 grid partially filled with four differently coloured circles. Participants were asked to identify the missing-coloured circle marked with a red question mark. The relationship between coloured items adhered to a single rule: *each item could appear only once per row and once per column*. This rule allowed us to manipulate relational demands by selectively removing items from the grid (Figure 1B).

**Figure 1.**
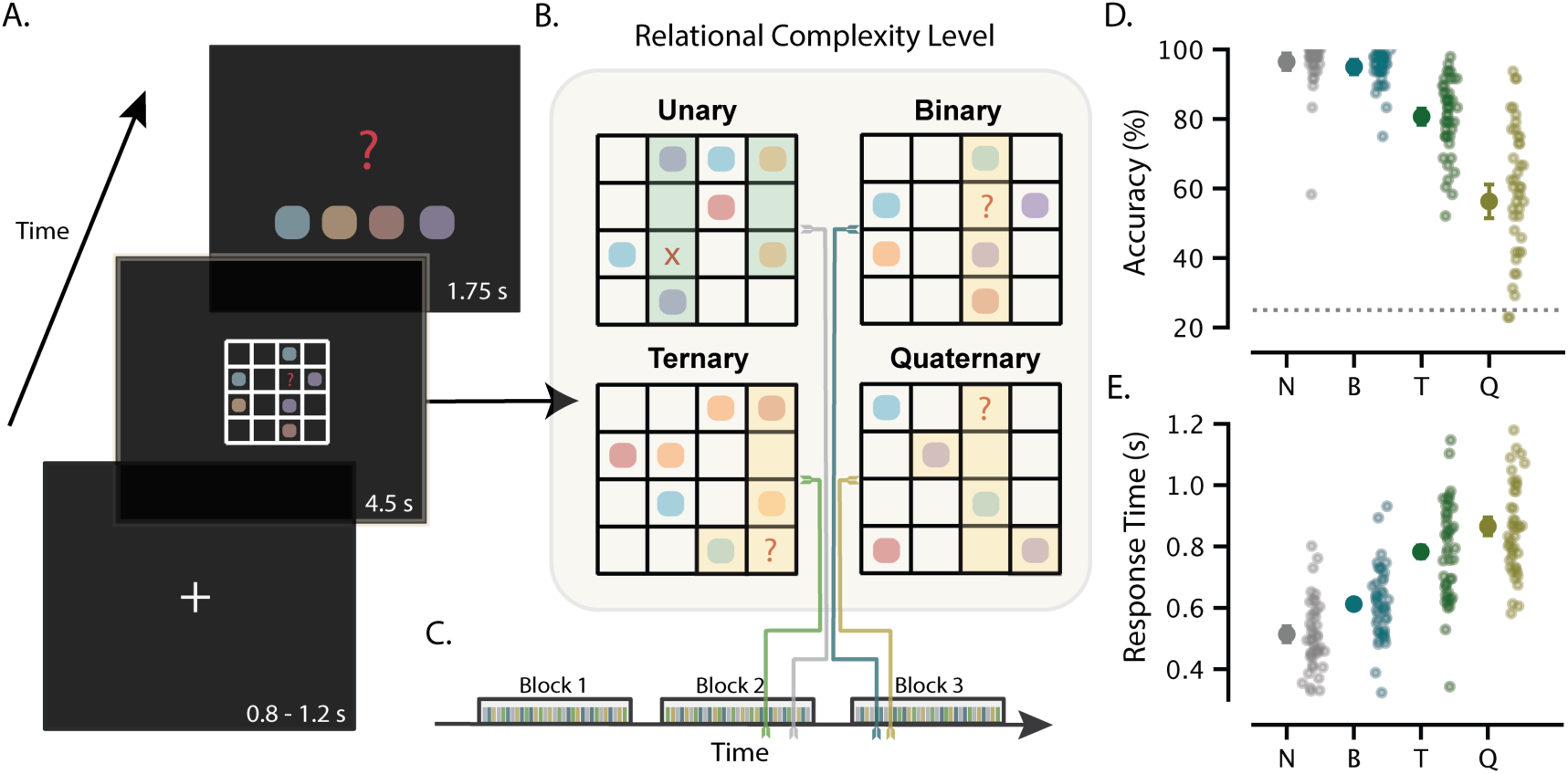
Latin Square Task adapted for concurrent EEG recording. (A) At the start of each trial, a white fixation cross appeared for a variable duration (0.8 – 1.2 seconds), then a Latin-Square puzzle was presented for a fixed 4.5 seconds, followed by a 1.75-second response screen. (B) 192 puzzles presented different levels of relational complexity (RC): unary/null, binary, ternary, quaternary. (C) 48 trials per RC level were presented across three event-related blocks, separated by a break determined by the participant. Each colour line depicts puzzles at a particular complexity level (grey, null; blue, binary; green, ternary; gold, quaternary) presented in pseudo-randomised order. (D) Subject-specific and group accuracy, with chance level (25%) depicted as the grey dotted line and (E) response time across RC levels: null (N), binary (B), ternary (T) and quaternary (Q). For each RC condition, the mean and within-subject 95% confidence interval are reported; points show individual performance (n = 45).

Our task design included four distinct levels of complexity, classified by the minimum number of chunked relations required to deduce the missing item (Figure 1B). Control problems (“unary” condition) presented unsolvable puzzles that violated the abstraction rule by containing duplicate items within the same row or column. These problems served as a baseline (“null” complexity) for rule comprehension. *Binary* problems required participants to combine items within a single row or column, using an internal representation of the complete set, to determine which of the four possible items could occupy that position. Increasing the requirement to coordinate information across both a row and a column yielded *ternary* problems. Finally, problems requiring participants to combine information across multiple rows and columns simultaneously were classified as *quaternary*, the highest level of relational complexity in the task.

As in previous studies that adopted the LST and related RC tasks, participants’ performance on our task declined as relational integration demands increased. Across the four complexity levels, performance accuracy declined as RC increased (Figure 1D; F(1.43, 62.99) = 111.54, p < 0.001, η_p_² = 0.717). Conversely, response times (Figure 1E) showed a progressive increase with complexity (Figure 1E; F(2.20, 96.64) = 144.74, p < 0.001, η_p_² = 0.767).

For the remainder of the study, all neurorecording analyses were restricted to correct trials (and related response times) to isolate neural activity that supports successful relational integration. Participants with insufficient artifact-free trials to yield reliable estimates were excluded (n = 11; see Methods and Figure S1), leaving a final sample of n = 34. Behavioural effects of relational complexity were preserved in this subgroup.

### Frontal and parietal activity showed dissociable patterns of activity as a function of increased relational integration demands

Our first goal was to characterise how local neural activity in frontal and parietal cortices mapped to different RC levels during the relational integration window (2 – 4.2 seconds; ^15^). We computed EEG signal power as a simple proxy for local synchronisation within these regional circuits. We started by visualising the time-frequency decomposition across participants, normalised to the pre-stimulus baseline (the inter-trial fixation cross, Figure 1A). Synchronisation is thus represented as a percentage change in power following puzzle onset (representative results from the left hemisphere shown in Figure 2A). Specifically, we plotted broadband (2 – 45 Hz) and band-specific (theta, alpha, beta, Methods) changes in signal power over time, highlighting the predefined relation integration window.

**Figure 2.**
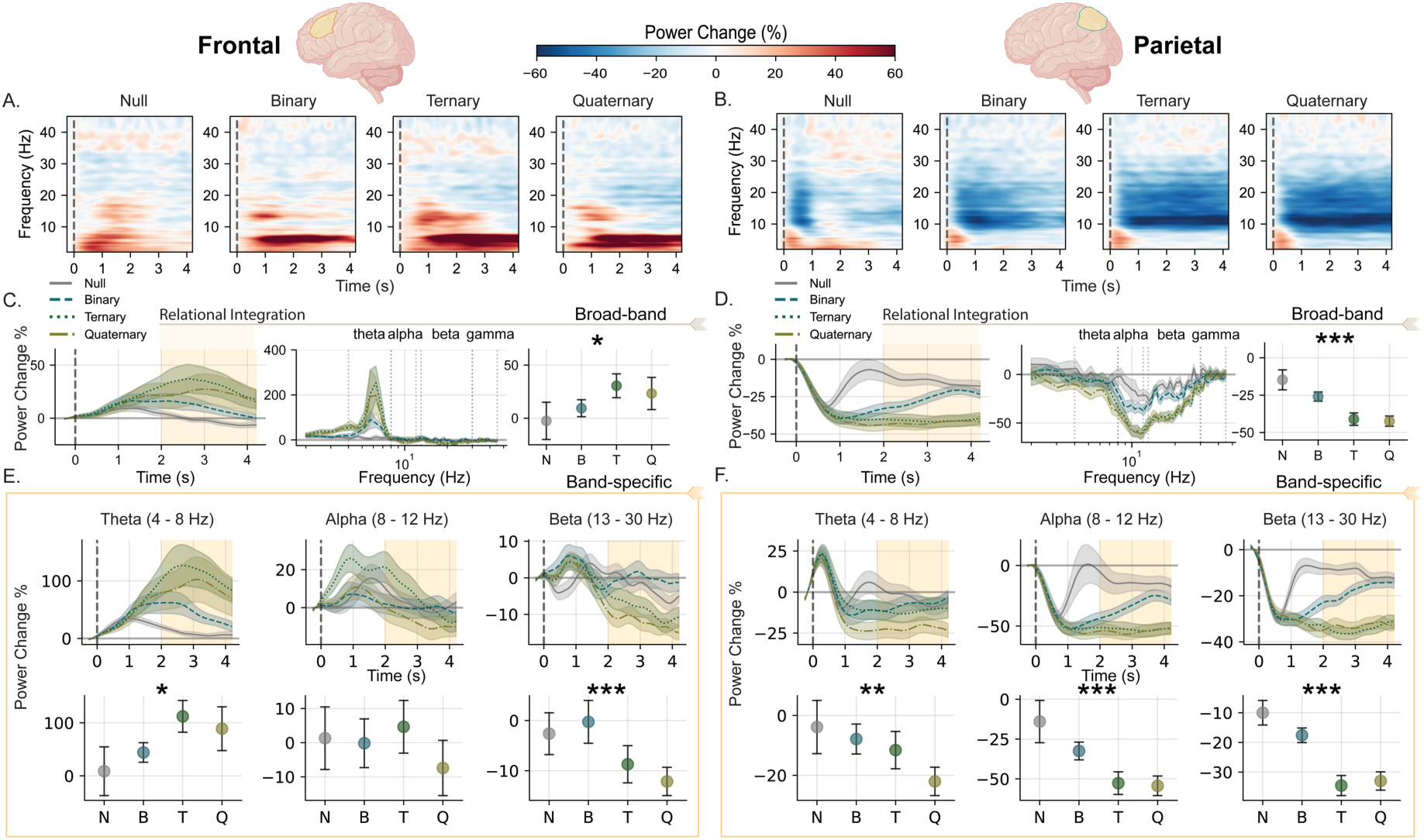
Changes in frontal and parietal cortical power as a function of increased relational complexity. Group-level spectral power dynamics across frontal (left panels) and parietal (right panels) regions during relational processing (n = 34). (A, B) Time-frequency representations (2-45 Hz) showing percentage power change from pre-stimulus baseline (−0.3 to −0.05 s) across four complexity conditions (null, binary, ternary, quaternary). Warm colours indicate increases in EEG signal power; cool colours indicate decreases in power relative to baseline. (C, D) Broadband power (2-45 Hz) time series across the puzzle stimulus, with shaded regions representing standard error of the mean. Baseline normalised power spectra (centre) extracted during the relational integration window (2-4.2 s, highlighted in beige ^15^). Interval plots (right) display condition means with within-subject 95% confidence intervals. (E, F) Band-specific time-series for theta (4-8 Hz), alpha (8-12 Hz), and beta (13-30 Hz) frequencies, with mean EEG signal power during the relational integration window below. Asterisks indicate significant differences between conditions (*p_FDR_ < 0.05, **p_FDR_ < 0.01, ***p_FDR_ < 0.001). Left hemisphere data shown.

The relative change in group-level power across complexity levels suggested opposing changes in signal power in left frontal (Figure 2A) and parietal (Figure 2B) cortices. Results showed that frontal mean broadband power increased with relational demands (F(1.20, 39.71) = 4.94, p_FDR_ = 0.039, η_p_^2^ = 0.13; Figure 2C). Conversely, the parietal mean broadband power decreased with increasing relational demands (F(1.64, 54.08) = 34.70, p_FDR_ < 0.001, η_p_^2^ = 0.51, Figure 2D). A similar trend was observed in the right-hemisphere frontal and parietal areas (Figure S2). Inspection of the relative change in the power spectral density (Figure 2C and D, middle panels) suggested that regional changes in broadband power were driven by discrete band-specific activity.

Analysis of narrowband EEG showed that broadband increases in left frontal regions were driven by theta power (Figure 2E, *left;* F(1.23, 41.69) = 7.31, p_FDR_ = 0.008, η_p_^2^ = 0.181), where beta-band power (Figure 2E, *right*) decreased with complexity (F(2.60, 85.95) = 10.54, p_FDR_ < 0.001, η_p_^2^ = 0.242). Frontal alpha power did not significantly differ. Similar results were observed for the right hemisphere (Figure S2).

In the parietal cortex, frequency-resolved power decreased with relational load (Figure 2F). The largest reduction in power occurred in the alpha (F(1.34, 44.28) = 20.24, p_FDR_ < 0.001, η_p_^2^ = 0.380) and beta bands (F(2.39, 75.68) = 63.17, p_FDR_ < 0.001, η_p_^2^ = 0.657), with comparable reductions in the right parietal cortex alpha (F(1.52, 50.00) = 14.30, p_FDR_ < 0.001, η_p_^2^ = 0.302) and beta bands (F(2.28, 75.32) = 32.76, p_FDR_ < 0.001, η_p_^2^ = 0.498; Figure S2). Theta power by contrast showed hemispheric asymmetry. It decreased with relational load in the left parietal cortex (F(1.97, 64.99) = 5.87, p_FDR_ = 0.008, η_p_^2^ = 0.151) but did not differ across conditions in the right parietal cortex (F(2.06, 68.04) = 0.255, p_FDR_ = 0.782, η_p_^2^ = 0.008).

We next asked whether these effects reflect stimulus-locked processes that are consistent across puzzles or induced activity that varies from trial to trial. We predicted that relational complexity effects would be driven by induced rather than phase-locked activity processing ^36–40^. To test this hypothesis, we decomposed the total spectral power presented in the above analysis into evoked (phase-locked) and induced (non-phase-locked) components and repeated the broadband and band-specific analyses across all four RC puzzle conditions (Methods). Results showed that the need to bind relations at various RC levels was associated with non-phase-locked activity during the puzzle presentation (Figure S3 and S4, and related text). In the late processing window, the signal was dominated by induced activity, with minimal evoked contribution. Accordingly, subsequent analyses were conducted on total power without further decomposition, consistent with prior work ^21,41,42^.

A concern in prior work is that baseline normalisation may distort low-frequency components of the spectrum (<10 Hz) ^43^. Moreover, broadband and band-specific results could be driven by mechanisms underlying the 1/f-like aperiodic and band-specific periodic oscillations in the spectrum. To address this, we decomposed the untransformed post-stimulus spectra into aperiodic and periodic components during the relation integration window (Figure S5 and related text). Results from this analysis confirmed that the observed band-specific effects reflect genuine oscillatory modulations rather than artefacts of normalisation or global spectral shifts. These results also revealed a regionally dissociable pattern in aperiodic activity, with frontal regions showing increases in offset and slope, and parietal regions showing the opposite trend (Figure S5). Analyses of periodic activity further showed that frequency-specific modulations occur alongside these broadband changes. These results suggest that aperiodic and oscillatory dynamics jointly contribute to complexity-related neural processing, but with distinct regional profiles.

### Band-specific frontoparietal activity and phase synchronisation support behavioural performance across relational demands

Building on the region- and band-specific power effects described above, we next asked whether reasoning behaviour was linked to changes in FPN synchronisation, as observed in fMRI studies ^4,5,44^. For each participant, we calculated change scores from lower to higher complexity and tested whether changes in FPN synchronisation statistically mediated the relationship between changes in nodal power and changes in performance (Figure 3). We modelled changes relative to the highest level of complexity (Quaternary), contrasting Quaternary with Binary (Model 1) and Quaternary with Ternary (Model 2). This approach isolates neural and behavioural changes associated with increasing relational demands as task complexity approaches its upper bound. We found no evidence that changes in FPN synchronisation mediated the relationship between nodal power and behavioural performance (Figure 3; Figure S6). However, examination of the individual pathways suggested that the relationships between FPN activity and behavioural performance differed according to both relational complexity and frequency band.

**Figure 3.**
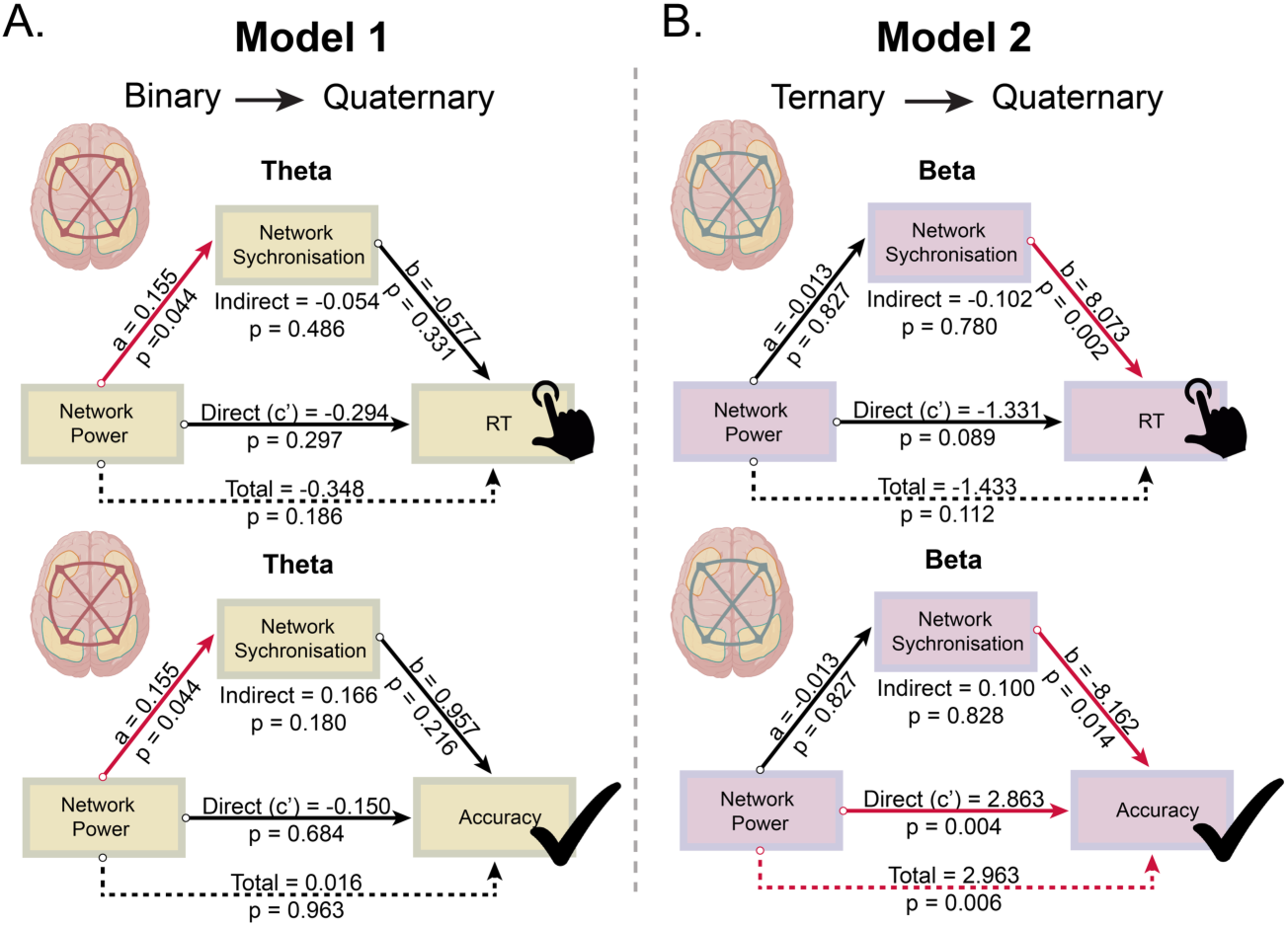
Relationship between changes in frontoparietal network activity and behaviour as relational complexity demands increase. Mediation models examine how changes in theta and beta FPN power across nodes influence behaviour (response time, *top*; accuracy, *bottom*) through changes in network synchronisation (debiased weighted Phase Lag Index; Methods) for (A) Model 1: the transition between low to high (Quaternary *minus* Binary) levels of complexity and (B) Model 2: the transition between medium to high (Quaternary *minus* Ternary) complexity. Mediation analysis performed using a bias-corrected non-parametric bootstrap method (95%, CI 5000 iterations). Path coefficients and p-values are reported for each pathway (black arrows). Significant individual regression paths are indicated by red arrows (p < 0.05). Because response time and accuracy reflect performance in opposite directions, the signs of the coefficients should be interpreted separately for each outcome. For response time, a positive coefficient indicates that greater complexity-related increases in power or synchronisation are associated with slower responses, whereas a negative coefficient indicates faster responses. For accuracy, a positive coefficient indicates that greater complexity-related increases in power or synchronisation are associated with higher accuracy, whereas a negative coefficient indicates lower accuracy.

For the largest increase in relational complexity (Quaternary–Binary; Model 1, Figure 3A), theta-band activity showed coordinated increases in frontal–parietal power and network synchronisation. However, these changes were not associated with behavioural performance, suggesting that while FPN theta dynamics are sensitive to relational demands, they do not directly account for individual differences in task performance. In contrast, the more incremental increase in complexity (Quaternary–Ternary; Model 2, Figure 3B) revealed a behaviourally relevant dissociation within the FPN beta activity. Increases in beta-band inter-regional synchronisation were associated with longer response times and reduced accuracy, whereas increases in nodal beta power showed the opposite pattern, tending to predict faster and more accurate performance. This suggests that at higher RC levels, behavioural performance results from a trade-off between local recruitment and inter-regional synchronisation, rather than simply increased network engagement. Across theta and beta band models, beta-band FPN activity emerged as a sensitive marker of relational integration performance; all models are reported in Figure S6.

### Distinct complexity-related corticothalamic gain modulation in frontal and parietal cortices

Given the observed changes in frontal and parietal activity and cross-regional synchronisation with increasing relational complexity, we next examined potential circuit-level mechanisms using a corticothalamic neural field model (CTM) applied to the source-reconstructed EEG spectra (Methods, Figure 4A). Within this framework, scalp EEG activity reflects the balance between the activity of cortical excitatory pyramidal neurons and inhibitory interneurons, while thalamic contributions arise from interactions between excitatory activity in the relay nuclei and inhibitory activity in the reticular nucleus ^29,45^. We interpret the fitted parameters in three families: local cortical gains, corticothalamic loop gains, and temporal filtering parameters. Local cortical gains index the balance of recurrent excitation and inhibition within each cortical region. Corticothalamic and intrathalamic gain index estimated feedback through relay and reticular thalamic populations. Synaptodendritic parameters index the duration of postsynaptic integration. Because we did not observe significant lateralisation effects in the parameter estimates (p_FDR_ > 0.05), we averaged the frontal and parietal model parameters across both hemispheres for subsequent analyses.

**Figure 4.**
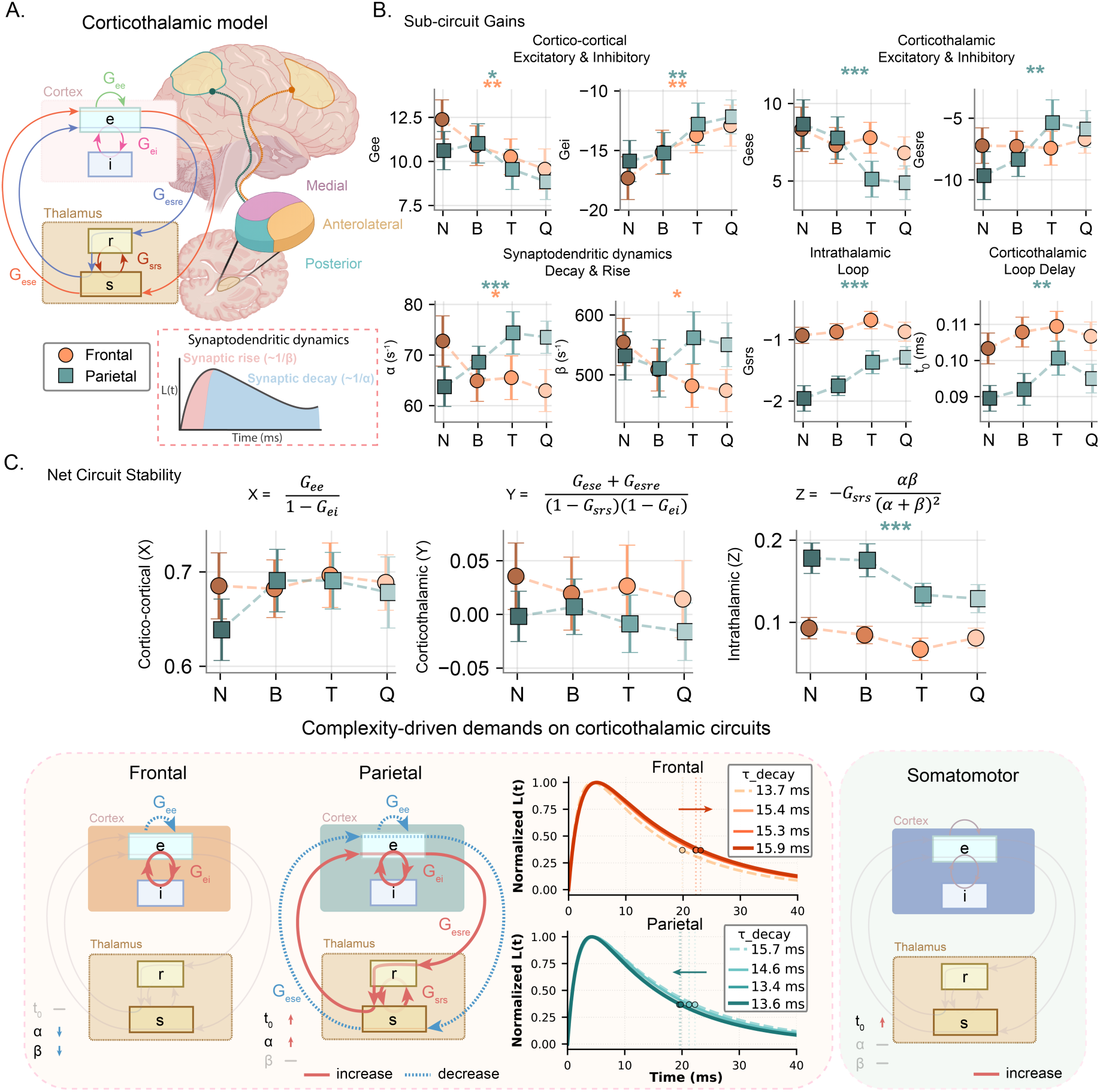
Model-derived engagement of the cortico-thalamic system during the encoding of relational complexity. (A) Schematic of the corticothalamic model (CTM): cortical excitatory (Gee) and inhibitory (Gei) populations, inhibitory thalamic reticular (Gsrs) and excitatory thalamic relay neural populations (Gese), corticothalamic inhibitory loop gain (Gesre) and their connections to frontal (orange) and parietal (teal) regions. Synaptic decay *α* and rise *β* describe the synaptodendritic dynamics. (B) CTM parameters across increasing relational complexity levels (N = null, B = binary, T = ternary, Q = quaternary relations). Colour fades from dark to light as complexity progresses. Top panels show cortical and corticothalamic excitatory and inhibitory gains; bottom panels show synaptodendritic rise and decay, intrathalamic loop gain, and corticothalamic loop delay. (C) Net cortical (left), corticothalamic (middle), and intrathalamic (right) gain across complexity levels. Bottom: Schematic summary illustrating differential corticothalamic circuit reorganisation as a function of increasing complexity for frontal (left) and parietal (right) regions. Significant main complexity effect for each region *p_FDR_ < 0.05, **p_FDR_ <0.01, ***p_FDR_ <0.001. Data show mean ± 95% within-subject confidence intervals.

Analysis of the circuit subcomponents revealed a coordinated modulation of cortical excitatory and inhibitory gains as relational complexity increased. Cortical excitability gain (G_ee,_ capturing recurrent excitation within the cortex) showed a progressive reduction with increasing complexity in both frontal (F(3, 99) = 4.925, p_FDR_ = 0.007, η_p_² = 0.130) and parietal (F(3, 99) = 3.471, p_FDR_ = 0.035, η_p_² = 0.095) regions (Figure 4B, top left). Conversely, cortical inhibitory gain (G_ei_, local cortical feedback inhibition) was weakened with complexity in both frontal (F(3, 99) = 5.262, p_FDR_ = 0.006, η_p_² = 0.138) and parietal (F(3, 99) = 4.887, p_FDR_ = 0.007, η_p_² = 0.129) regions (Figure 4B, top left).

Corticothalamic coupling showed a more region-specific pattern. In the parietal cortex, excitatory corticothalamic loop gain (G_ese,_ capturing the direct positive feedback loop between the cortex and the thalamic relay nuclei) decreased with increasing complexity (F(3, 99) = 8.696, p_FDR_ < 0.001, η_p_² = 0.209). Conversely, excitatory corticothalamic loops had no significant effect on frontal activity (F(3, 99) = 1.301, p_FDR_ = 0.360, η_p_² = 0.038). Parietal corticothalamic inhibitory loop gain (G_esre_, capturing indirect inhibitory feedback via the thalamic reticular nucleus) was attenuated by relational demands (F(3, 99) = 6.044, p_FDR_ = 0.004, η_p_² = 0.155), with no corresponding change in the frontal cortex (F(3, 99) = 0.242, p_FDR_ = 0.908, η_p_² = 0.007; Figure 4B, top right). Parietal corticothalamic loop delay (t_0_) increased as a function of relational complexity (t_0_: F(3, 99) = 5.447, p_FDR_ = 0.006, η_p_² = 0.142), indicating slower corticothalamic signal propagation. Within the thalamus, intrathalamic loop gain (G_srs_, reticular gating of relay neurons, capturing local thalamic inhibitory feedback) was weakened as relational demands increased (F(3, 99) = 12.477, p_FDR_ < 0.001, η_p_² = 0.274). No significant complexity-related changes were observed for frontal intrathalamic loop gain (F(3, 99) = 2.463, p_FDR_ = 0.105, η_p_² = 0.069) or corticothalamic delay (t_0_: F(3, 99) = 1.633, p_FDR_ = 0.257, η_p_² = 0.047; Figure 4B, bottom right).

Synaptodendritic temporal parameters, which govern the temporal filtering of synaptic inputs throughout the corticothalamic system, exhibited opposing regional adaptations in postsynaptic rise and decay with increasing relational complexity (Figure 4B, bottom left). In parietal regions, the synaptic decay rate constant (*α*, the rate at which postsynaptic potentials decay at the soma) increased with complexity, suggesting a shorter integration window. In contrast, synaptic rise (*β*, the rate of postsynaptic potential onset) remained unchanged (decay: F(3, 99) = 7.667, p_FDR_ < 0.001, η_p_² = 0.189; rise: F(3, 99) = 1.168, p_FDR_ = 0.398, η_p_² = 0.034). Conversely, results in frontal regions showed decreases in both synaptic decay and rise rate constants with increasing relational complexity (decay: F(3, 99) = 3.980, p_FDR_ = 0.020, η_p_² = 0.108; rise: F(3, 99) = 4.136, p_FDR_ = 0.018, η_p_² = 0.111). Together, these opposing synaptodendritic tuning patterns demonstrate a regional dissociation in the temporal integration of post-synaptic potentials. Parietal regions exhibit faster filtering of synaptic inputs, whereas frontal regions support more sustained synaptic integration as relational demands increase.

We also examined a reduced model in which circuit stability is characterised by the net gains in cortico–cortical (X), corticothalamic (Y), and intrathalamic (Z) loop feedback (Methods Equations 13-15, Figure 4C). Neither cortico–cortical nor corticothalamic loops showed a significant net gain with relational demands (all p_FDR_ > 0.05), suggesting that increasing relational integration demands were not accompanied by broad circuit-wide reorganisation at the cortical or corticothalamic level. Instead, stability was maintained through parameter-specific gain adjustments that balanced excitatory and inhibitory dynamics within the circuit (see above, Figure 4B). Intrathalamic net gain, however, exhibited a significant complexity-dependent decrease in parietal regions (F(3, 99) = 9.362, p_FDR_ < 0.001, η_p_² = 0.221). Frontal regions exhibited a similar but non-significant reduction in intrathalamic gain (F(3, 99) = 3.105, p_FDR_ = 0.051, η_p_² = 0.086).

As a control, we also examined a cortical set of regions not thought to be primarily involved in complexity representations: the somatomotor network ^5,15^. Moreover, higher-order cognition appears to engage medial and posterior thalamic neurons, while sensory-motor activity preferentially activates the anterolateral portion (Figure 4A; ^26^). No significant changes in sub-circuit or net corticothalamic gain parameters were observed across complexity levels (all p_FDR_ > 0.05; Figure S7). The only effect was an increase in corticothalamic loop delay (F(3, 99) = 4.743, p_FDR_ = 0.044, η_p_² = 0.125; summarised in Figure 4), suggesting that the prolonged delay observed during relational processing may reflect a more general task-induced slowing in thalamo-cortical coupling. These control modelling findings highlight that changes in gain and stability observed in frontoparietal regions reflect regionally specific adaptations associated with relational integration.

## DISCUSSION

Humans can integrate multiple relations, enabling reasoning beyond simple associations ^46^. Here, we showed that frontal and parietal regions may support this process through distinct neural mechanisms. As complexity increased, frontal and parietal regions exhibited distinct load-dependent changes in oscillatory power and synchronisation. Biophysical modelling extended these results by revealing a circuit-level division. The frontal cortex supported longer-sustained neural integration by modulating the local excitatory-inhibitory balance, while parietal regions reorganised thalamocortical loops to accommodate relational integration demands. Together, these findings reframe frontoparietal interactions not as a uniform recruitment of shared resources, but as regionally differentiated computational strategies in which synchronisation is necessary but not sufficient for problem-solving, and in which excitatory–inhibitory rebalancing, rather than gain amplification, is likely key to supporting relational reasoning.

At the regional level, increasing relational complexity was associated with opposing shifts in broadband power at the frontal and parietal cortices. Spectral parameterisation indicated that these effects were driven by region-specific changes in aperiodic activity, alongside selective recruitment of periodic oscillations. Frontal regions showed an upward shift in aperiodic activity with increasing demand, whereas parietal regions showed a downward shift, indicating divergent adaptations across the frontoparietal system. Aperiodic spectral features have recently emerged as a sensitive, domain-general index of cognitive demand across tasks, including serial subtraction, working memory, task switching, and interference paradigms ^20,22^. Extending this literature, the present findings demonstrate that aperiodic adaptations during relational integration are not uniform but regionally differentiated, suggesting distinct computational roles within the frontoparietal system.

Superimposed on these broadband shifts were dissociable band-limited effects. Increased relational complexity was characterised in the frontal cortex by enhanced theta power and attenuated beta power, whereas the parietal cortex showed reductions in both alpha and beta power. This spectral profile differs from prior representational analyses of the same time window, which showed that behaviourally relevant representations of relational complexity are primarily encoded in frontoparietal alpha and beta bands, consistent with dynamic redistribution of representational weight as demands increase ^15,16^. Together, these findings suggest that stable encoding and online construction of relations rely on partly separable mechanisms: alpha and beta support representational stability, while frontal theta increases with demand, reflecting dynamic coordination for binding multiple elements. This interpretation supports functional accounts linking lateral frontal theta to top-down control and adaptive network coordination during cognitively demanding operations ^47–50^. Accordingly, Lu *et al.* ^22^ demonstrated frontal theta recruitment across distinct cognitive control paradigms that did not explicitly manipulate relational integration. The present findings extend previous work by showing that lateral frontal theta recruitment is sensitive to the problem’s complexity, suggesting that integrative binding demands may be a specific computational driver of control-related theta dynamics.

Consistent with prior work, increasing relational demands were accompanied by reductions in parietal alpha and beta activity, often linked to higher cognitive load and active processing ^21,41,42^. However, this pattern is not universal; some studies report increases in these bands with task difficulty ^22^. This discrepancy may reflect differences in task structure. Our multi-level complexity manipulation revealed nonlinear neural recruitment that simpler dual-level contrasts may have missed. More broadly, these findings suggest that spectral modulation depends on how information is used. Beta attenuation may reflect a shift from stable maintenance to flexible processing, indicating that relational integration preferentially engages dynamic updating rather than sustained representations.

Our mediation analysis identified that covarying frontoparietal theta and beta power and inter-regional synchronisation differentiate discrete levels in relational processing demands, building on previous fMRI research showing increased integration during relational tasks ^2,4,5,7,51,52^. Beta activity was behaviourally relevant, but greater synchrony was not always beneficial. Excessive network engagement predicted poorer performance at higher complexity. This may indicate that efficient coordination, rather than maximal activation, supports reasoning. Beta effects were linked to transitions to the highest complexity, consistent with a role in large-scale coordination. These findings highlight the importance of interhemispheric and long-range synchronisation for higher-order relational processing, with disruptions in such coordination likely constraining the formation of complex relational abstractions ^44,53,54^.

Complementing the empirical results, the CTM modelling results suggest that increasing complexity was associated with reduced excitatory and inhibitory cortical gains, maintaining balance. In addition, relational encoding demands induced regionally dissociable adjustments in synaptodendritic temporal dynamics. Parietal cortex exhibited faster synaptic decay as complexity increased, consistent with a narrowing of the temporal integration window. Conversely, the frontal cortex showed concurrent slowing of both rise and decay kinetics, supporting more sustained integration ^29,30,55^. These opposing adjustments suggest complementary temporal regimes, in which faster parietal decay may enhance temporal precision and rapid updating of relational representations, while prolonged frontal integration may support sustained maintenance and manipulation over extended time scales.

Cortico-cortical and corticothalamic net gain remained largely unchanged across complexity, indicating that stability is maintained through compensatory local adjustments rather than large-scale reconfiguration ^30^. The exception was a selective reduction in intra-thalamic gain in parietal circuits, suggesting downregulation of thalamic feedback specific to higher-order processing. Together, these findings suggest that relational complexity is supported by targeted local modulation in local gain, synaptic timing, and thalamic feedback while preserving global circuit stability; a pattern that may generalise across other cognitive demands and domains ^20,22,42,56,57^.

Several considerations should be kept in mind when interpreting these findings. Although source reconstruction was used to estimate cortical activity, the spatial resolution of the present measurements remains constrained by scalp EEG. Moreover, the corticothalamic parameters were inferred by fitting a biophysical neural field model to source-reconstructed cortical spectra and should therefore be interpreted as model-derived estimates of plausible circuit mechanisms rather than direct physiological measurements. The use of the Latin Square Task was motivated in part by prior high-field fMRI evidence implicating the thalamus in relational complexity^26^, but confirmation of the specific thalamic mechanisms inferred here will require complementary approaches, including invasive recordings or targeted causal perturbation. Distinct parameter configurations may also produce partially overlapping spectral effects. Finally, although relational complexity is considered a domain-general property of reasoning^46^, replication across other task paradigms will be necessary to determine whether these circuit adaptations generalise beyond the present task. Our focus on predefined frontal and parietal regions within an independently established relational-integration window constrained the analysis space in a hypothesis-driven manner, but may have overlooked relevant dynamics outside these spatial and temporal boundaries.

In summary, the current work offers a novel multiscale perspective on how fronto-parietal-thalamic neural dynamics at regional, inter-regional, and circuit levels adapt to increasing relational complexity. The results show that relational complexity is resolved by region-specific modulations in neural activity that maintain the global system’s stability.

## METHODS

### Participants

Forty-seven participants (mean age = 25; age range = 19 – 33; 62% female) were recruited from the Brisbane Metropolitan area. The sample was largely composed of university students or individuals with higher education. Ethics approval was granted by the QIMR Berghofer Research Ethics Committee (P3644). Participants provided written informed consent and were eligible if they were aged between 18 and 35 years and had no previous reported history of mental health or neurological disorder.

The number of participants was based on an a priori power analysis performed using G*Power (version 3.1) to estimate the statistical power required to detect a behavioural effect between two dependent means (matched pairs). An alpha of 0.05 and a desired statistical power of 0.8 required a minimum sample size of 34 participants to detect medium-to-large effects (d > 0.5). Additional participants (n=13) were recruited to account for a 15% attrition rate, EEG acquisition errors, and exclusions during preprocessing (see section EEG preprocessing). Participation in this study was voluntary, and participants were reimbursed $20 AUD an hour for their time.

Of the 47 recruited participants, 2 were excluded due to poor data acquisition, leaving 45 participants for behavioural analyses; a further 11 were excluded from EEG analyses for failing to meet the minimum correct trial threshold (≥20 trials per condition; Figure S1), yielding a final EEG analysis sample of 34 participants.

### Latin Square Task

The LST is a nonverbal relational reasoning paradigm that systematically varies processing demands by manipulating the number of relations that must be interrelated simultaneously (Birney et al., 2006). Participants are presented with a four-by-four grid containing coloured circles and a target question mark (Figure 1). Participants must solve for the question mark by applying the rule that an item (in this case, one of four colours) can appear only once in a row or column. Task complexity is manipulated across four levels based on the number of relations that must be processed simultaneously to determine the solution. Binary relations require processing along a single dimension (either a row or a column). Ternary relations require integrating information across a single row and a single column. Quaternary relations demand simultaneous processing across multiple rows and columns. Control puzzles were designed as unary relations that intentionally violate the task’s underlying logic and are unsolvable, as indicated by the ‘x’ response option. For example, these puzzles might contain two purple elements in the same column, violating the fundamental Latin Square constraint that each element appears exactly once in each row and column. Due to this low complexity, unary relational operations are indistinguishable from simple associations and thus serve as an appropriate control condition against which to compare more complex relational reasoning demands.

The original LST has been previously validated and adapted for fMRI neuroimaging acquisition ^13^. Here we further modified the design for EEG ^15^. To generate sufficient trials using the original twelve unique puzzle designs per complexity level, each original puzzle was rotated by 90 degrees three times and the item positions were exchanged. This preserved the underlying relational topology while creating novel visual configurations. The experiment comprised of 48 trials per complexity level (12 unique puzzles × 4 transformations). Trials were pseudo-randomised to prevent sequential presentation of the same complexity level (i.e., to avoid the repetition of two binary problems). We further modified the stimulus presentation time, shortening the puzzle presentation to a 4.5-second window followed by a 1.75-second response window.

### EEG acquisition and procedure

EEG data were recorded using a 64-channel ANT-Neuro system (eego sports) with electrodes positioned according to the international 10-20 system. Signals were digitised at 2000 Hz with electrode impedances maintained below 10 kΩ using conductive gel. Stimuli were presented on a 24-inch BenQ monitor using PsychoPy (v2020.2.10; Peirce et al., 2019). To minimise eye movements, the puzzle grid was sized to fit within the participants’ central field of view; in addition, participants were instructed to minimise their eye movements.

Following an explanation of the task, participants began by completing 10 practice trials with numerical stimuli (1-4). The experimenter provided feedback to the participants to ensure they understood the task. Puzzles were assigned arbitrary names (Bliss, Argon, York, Unsolvable) to avoid framing the problems as easy or hard based on the RC classification level (Binary, Ternary, Quaternary, and Null conditions). During the response screen, participants use a computer keyboard to press keys 1, 2 (left hand) or 9, 0 (right hand) corresponding to one of the four items (Figure 1). For the main experiment, stimuli were changed to colours, and participants completed 192 trials divided across three blocks without performance feedback.

### EEG preprocessing

EEG data were preprocessed offline using custom scripts adopting EEGLAB functions ^58^ in MATLAB (v2019b). The raw continuous data underwent high-pass finite impulse response (FIR) filtering at 0.5 Hz to remove low-frequency drifts. Line noise centred at 50 Hz was eliminated using a notch filter spanning 47.5 - 52.5 Hz, followed by a 97 Hz low-pass filter to attenuate harmonic components. Bad channels, identified through visual inspection, were excluded before re-referencing to the average of all remaining channels, excluding the M1, M2, and EOG electrodes. The data were then downsampled to 250 Hz.

Trial epochs were generated using the stimulus (puzzle) onset as a marker, resulting in a −400 ms pre-stimulus and 4500 ms post-stimulus window. Trials heavily contaminated by muscle artefacts were identified through visual inspection and rejected. Independent component analysis (ICA) was then applied to identify and remove the remaining non-neural activity, including eye blinks, cardiac activity, and residual muscle artefacts. Following ICA-based artefact rejection, channels excluded during preprocessing were reconstructed using spherical spline interpolation. Finally, the signal underwent band-pass filtering within the EEG sensitive range of 1-45 Hz.

Only correct trials were included in the final analysis, excluding incorrect or missing responses to aid in result interpretation. Participants were included in subsequent analyses if they met a minimum criterion of 20 correct trials per condition, a trial threshold similar to that used in other EEG task studies ^21,43^. To validate this criterion, we conducted a reliability analysis determining the number of trials needed to obtain stable power measures across frequency bands and regions of interest ^37^ (Figure S1). For each participant, we created a task-average template by concatenating power activity across all conditions, then iteratively sampled increasing numbers of correct trials and correlated both total and band-specific power with the grand average. Reliability estimates (r-values) were averaged across subjects to assess the convergence of broad and band-specific signal power as a function of trial number. To address unbalanced trial numbers between conditions resulting from using only correct trials and artifact rejection, we randomly downsampled trials in each condition to match the condition with the lowest trial count (typically the quaternary condition). Figure S1, displays the mean and inter-quartile range of the balanced trial numbers per condition for each region.

### LST Relational Integration Window

The temporal analysis window and ROIs were chosen based on a prior multimodal localisation study that identified when and where relational complexity representations emerge during puzzle solving of the Latin Square Task ^15^. That study combined representational similarity analysis of the present EEG dataset (n = 45) with an independent fMRI cohort (n = 40) and fMRI–EEG fusion analyses to localise the timing of RC representations. Across modalities, RC representations consistently emerged during the later stages of reasoning, with significant effects occurring 2.1 – 4.2 s after puzzle onset, the precise interval varying across frequency bands. Guided by these results, the analyses in the present study focus on a 2.0 – 4.2 s period after puzzle onset, termed the *relational integration window*. The window onset starts at 2.0 s to conservatively capture the earliest emergence of RC-related activity across analyses and to provide a common window across conditions and frequency bands. Importantly, this choice is independent of the analyses reported here. While the previous study characterised information contained within the multivariate representational geometry of puzzle states at the whole-brain sensor (EEG) and voxel (fMRI) level, the present study examines source-level spectral power and biophysical circuit parameters within predefined cortical regions.

### Source modelling and time-series extraction

Spatial deconvolution of scalp-level EEG sensor recordings was implemented using the M/EEG toolbox Brainstorm ^59^. The forward model was generated using the Boundary Element Method (BEM) ^60,61^ with the default ICBM152 anatomical template ^62^. Cortical sources were reconstructed using dynamical Statistical Parametric Mapping (dSPM) onto a cortical mesh containing 15,000 vertices (henceforth referred to as sources) ^63^. Source model dipole orientations were constrained normal to the cortex, and default depth weighting parameters (order = 0.5; maximal amount = 10) were applied to adjust for bias toward superficial sources. The dSPM normalisation process utilised noise covariance statistics derived from each participant’s first five minutes of resting-state recording. This noise covariance matrix was used to estimate noise levels at each source location, and the minimum norm estimates were then divided by these location-specific noise estimates. This normalisation procedure yielded unitless z-score maps, indicating the number of standard deviations by which each source’s activity exceeded the estimated noise level. For noise covariance regularisation, we used the diagonal noise covariance matrix to help stabilise the inverse solution, with the regularisation parameter signal-to-noise ratio kept at the default value of 3. Cortical parcels were drawn from the Schaefer parcellation ^64^ registered to the Brainstorm ICBM152 anatomical template to generate frontal and parietal masks corresponding to FPN regions ^59^. In addition we selected the somatomotor network (SomMat) as a control region. Together, this procedure yielded six bilateral ROIs (left/right frontal, parietal, somatomotor; masks available on GitHub shown in Figure S8). To reduce the dimensionality of the source space, source activity within each ROI was averaged, resulting in a single time series per trial, condition, and ROI for subsequent analyses.

### Time-frequency analysis

We performed temporal and spectral decomposition of the frontal and parietal time series using a set of complex Morlet wavelets, defined as complex sine waves tapered by a Gaussian ^65^. To minimise edge artifacts, we mirrored 0.6 seconds of each epoch time series at both the beginning and end before convolution. The wavelet frequencies ranged from 2 Hz to 45 Hz in 82 linear steps. The full width at half-maximum (FWHM) ranged from 1000 to 200 ms, increasing with wavelet peak frequency. This corresponded to a spectral FWHM range of 0.88 Hz (low-frequency wavelets) to 4.4 Hz (high-frequency wavelets). Following wavelet convolution of the concatenated mirrored epochs, we re-segmented the data into epochs, removing the mirrored segments.

Time-frequency maps were decomposed into induced (non-phase-locked) and evoked (phase-locked) components across participants and conditions ^37^. Total power was computed by calculating time-frequency maps for each trial before averaging to generate condition-specific maps. Induced power was extracted by subtracting the condition-averaged ERP from single-trial activity before generating condition-averaged maps. Evoked activity was obtained by subtracting the induced activity from the total activity.

Each participant’s averaged time-frequency maps (for each condition and power type) were normalised using event-related change from the pre-stimulus baseline window (−0.3 to −0.05 seconds). The post-stimulus activity can thus be interpreted as event-related synchronisation (ERS) or event-related desynchronisation (ERD). Time series of broadband and band-specific energy were created by averaging activity within specific frequency ranges in the time-frequency maps: broadband (2 – 45 Hz), theta (4 – 8 Hz), alpha (8 – 12 Hz), and beta (13 – 30 Hz). To quantify changes in spectral power, both the untransformed and baseline-normalised activity were temporally averaged within the previously established relational integration window (2.0 to 4.2 seconds post-stimulus onset) ^15^.

To account for aperiodic slope dynamics we modelled the neural power spectra to parameterize aperiodic and periodic task-induced activity during the post-stimulus relational encoding period using the spectral parameterization algorithm (version 2.0.0rc2; https://specparam-tools.github.io/) ^66^. Based on our data characteristics and prior work ^20,21^, we modelled the spectrum between 4 – 40 Hz, as frequencies below this range become difficult to model and higher frequencies introduce a bend in the power spectra. The algorithm parameters were set as follows: peak width limits (1, 8), maximum number of peaks (inf), minimum peak height (0.0), peak threshold (2.0), and aperiodic mode (’fixed’). From the model, we extracted two key aperiodic parameters: the exponent describing the steepness of the spectral slope and the offset capturing shifts in broadband power. To allow comparison with the unparameterised band-specific results the mean periodic band-specific power (relative power) was also extracted. Good model fits (R^2^ ≥ 0.85) were attained across all subjects, regions and conditions.

### FPN synchronisation

Phase synchronisation between ROI pairs was quantified using the debiased weighted Phase Lag Index (dwPLI; ^67^). The dwPLI measures the consistency of phase differences between neural signals by analysing the asymmetry of phase difference distributions. This approach specifically detects non-zero phase relationships and eliminates spurious correlations from volume conduction by utilising only the imaginary component of the cross-spectrum ^68^.

To construct the frontoparietal network, dwPLI values were calculated for each trial across six possible edge connections between four FPN ROIs: bilateral frontal, bilateral parietal, and four frontal-parietal connections (left frontal-left parietal, right frontal-right parietal, left frontal-right parietal, right frontal-left parietal). For each edge, dwPLI was calculated at frequencies between 4-30 Hz using the wavelet decomposition, then averaged within three band-specific frequency bands: theta (4-8 Hz), alpha (8-12 Hz), and beta (13-30 Hz). Values were further averaged across the relational integration window (2 – 4.2 s) to generate a single matrix representing the synchronisation during relational processing. This yielded band-specific connectivity matrices (4×4×136) representing phase synchronisation patterns between all ROI pairs for each subject (n=34) and conditions (null, binary, ternary, quaternary).

### Statistical Analysis

Task accuracy, response time and regional power were analysed using a one-way within-subject ANOVA to assess for a main RC effect implemented in JASP. Where a Mauchly’s test indicated that the assumption of sphericity was violated (p < 0.05) a Greenhouse-Geisser correction was applied. Normality was initially assessed using a Shapiro-Wilk test and inspection of the distribution. A log transformation was applied to the raw power values as the power spectra follows a log-normal distribution. The main effect of complexity from the power analysis was false discovery rate (FDR) corrected for multiple comparisons using the Benjamini-Hochberg method implemented in MATLAB to correct for the four ROIs, power components, and frequency bands ^69^. For the CTM modelling analysis, FDR correction was applied across ROIs and the number of fitted parameters.

### Mediation Analysis

As an exploratory analysis, we conducted mediation analyses ^70^ using the Python package Pingouin (version 0.5.4; ^71^) to evaluate the hypothesised pathway whereby changes in FPN synchronisation mediate the relationship between changes in regional power and behavioural performance. For each band-specific network (theta, beta) we examined two transitions in RC: the most extreme (Quaternary-Binary) and incremental (Quaternary-Ternary). We opted to start from binary as the null condition is an association rather than reasoning. Individual difference scores were calculated by subtracting the lower complexity condition from the higher complexity condition for each measure (power, network synchronization, response time, accuracy). Mediation models were constructed with region node power differences as the independent variable (X), network synchronization differences as the mediator (M), and behavioural differences (response time or accuracy) as the dependent variable (Y). Significance was assessed using a bias-corrected bootstrap method with 5000 iterations and 95% confidence intervals. For each model, we report the regression coefficients and p-values for: the effect of power on network synchronisation (path a), the effect of network synchronisation on behaviour controlling for power (path b), the direct effect of power on behaviour controlling for network synchronization (path c’), and the indirect effect through network synchronization (path a*b).

### Corticothalamic Modelling of the EEG spectra

We applied a well-established corticothalamic model (CTM) ^29,72^ to estimate the biophysical parameters underpinning the frontal, parietal, and somatomotor power spectra during the relational integration window. Parameter estimation was implemented using the BrainTrak software library ^28,32^. This modelling approach has successfully described oscillatory dynamics, gain modulation, and state-dependent changes in corticothalamic excitability across sleep, ageing, focal thalamic pathology, and large-scale neurological disruption ^28,31–35^.

The model applies neural field theory to describe interactions between four coupled cortical and thalamic neuronal populations: cortical excitatory pyramidal neurons (*e*) and cortical inhibitory interneurons (*i*), together with excitatory thalamic neurons in the specific relay nuclei (*s*) and inhibitory thalamic reticular neurons (*r*). Model parameters and limits are reported in SI Appendix, Table 1.

Each neuronal population *a* ∈ {*e*, *i*, *s*, *r*} is characterised by a mean cell-body firing rate *Q_a_*, and a mean somatic membrane potential *V_a_*. The relationship between membrane potential and firing rate is governed by a sigmoid function that reflects the threshold-like aggregate input–output properties of a neural population:

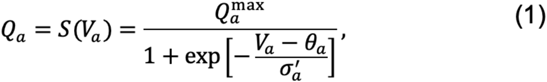

where, *Q^max^_a_* is the maximum firing rate, *θ_a_* is the mean firing threshold, and *σ^’^_a_ = σ_a_√3/π* characterises the spread of firing thresholds across the population. Hence, Equation 1 captures how an increase in net input to a population causes its collective output to shift nonlinearly from silence to near-maximal firing across the entire population.

The somatic membrane potential of each population (*a*) is determined by all synaptic inputs received from each presynaptic population *b*:

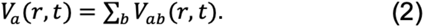

Each synaptic input component *V_ab_* is generated by the pulse flux arriving from presynaptic signals *φ_b_* after passing through a second-order synaptodendritic filter. This filter captures the time course of a typical postsynaptic potential (the rise and decay of synaptic current in the dendritic tree) and is expressed as:

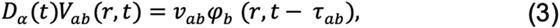

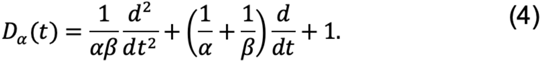

*D_α_*(*t*) acts as a temporal low-pass filter, where *α* is the inverse synaptodendritic decay time constant and *β* is the inverse rise time constant. Together, *α* and *β* control the temporal width of individual postsynaptic responses, where larger values imply faster synaptic kinetics and a narrower integration window, whereas smaller values imply slower, more sustained postsynaptic potentials. *ν_ab_* is the connection strength from population *b* to *a*, and *τ_ab_* is the mean axonal propagation delay between the two populations.

The mean axonal propagation (*φ_a_*) across the cortical sheet is modelled by a damped wave equation:

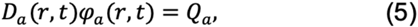

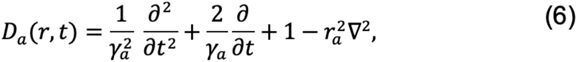

where *r_a_* is the characteristic axonal range of population *a,* and *γ_a_* = *v_a_*/*r_a_*is the temporal damping rate, where *v_a_* is the mean axonal conduction velocity. The Laplacian term *r^2^_a_∇^2^* encodes the spatial spread of activity, together capturing the dispersion relation governing oscillatory dynamics. The damped wave equation thus links the local firing rate *Q_a_* to the spatially distributed pulse flux *φ_a_*, producing the finite propagation speeds and spatial attenuation observed in cortical tissue. In the model, only excitatory axons are long enough to produce significant propagation effects. Wave propagation effects are assumed to be negligible for short-range axons of cortical inhibitory neurons and the thalamus, so the solution to Equations 5 and 6 is approximated by *φ_a_* = *S*(*V_a_*) for a = (*i, s, r*) ^73^. The model is further simplified by assuming that intracortical connections are random, with the number of connections proportional to the number of synapses and with symmetric connection strengths ^32^. Resulting in a model with eight independent connections.

### EEG power spectrum

Under linearisation around the steady state and transformation to the frequency-wavenumber domain (*k*, *ω*), the transfer function relating cortical excitatory activity *φ_e_* to a subcortical noise input *φ_n_* is:

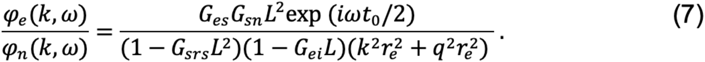

The numerator reflects the gain for subcortical inputs driving the cortex through the thalamic relay pathway, while the denominator captures resonances arising from corticothalamic and intracortical feedback loops. The parameter *t*_0_ is the total axonal conduction delay for a full corticothalamic round trip.

The gain parameters *G_ab_* = *ρ_a_ν_ab_* appearing throughout are defined as the product of the slope of the sigmoid function at its operating point, *ρ_a_*, and the synaptic strength *ν_ab_*. Gains involving an inhibitory postsynaptic population (*G_ei_*, *G_sr_*) are negative.

The dynamics obey a dispersion relation *k*^2^*r_e_*^2^ + *q*^2^*r_e_*^2^ = 0, where

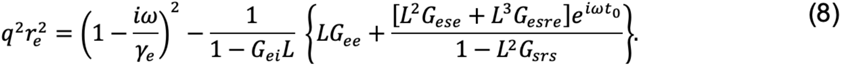

The first term on the right gives the free cortical dispersion in the absence of feedback; the remaining terms progressively modify this by intracortical excitation (*G_ee_*), corticothalamic feedback through the excitatory relay loop (*G_ese_*), and feedback through the longer corticothalamic–reticular inhibitory loop (*G_esre_*), all gated by intracortical inhibition (*G_ei_*) and inhibitory intrathalamic loop gain (*G_srs_*). Resonances in the power spectrum arise when *q*^2^ ≈ 0 (i.e., when the denominator of Equation 7 is small), reflecting instability in coupled feedback loops.

The synaptodendritic transfer function *L*(*ω*) is:

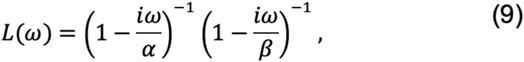

which behaves as a low-pass filter, producing the characteristic broadband 1/f-like scaling of the EEG spectrum.

### Obtaining the power spectrum

The modelled EEG power spectral density is obtained by summing the squared transfer function across all cortical spatial modes, weighted by a spatial filter function that reflects the finite aperture of scalp EEG electrodes relative to the spatial scale of cortical activity:

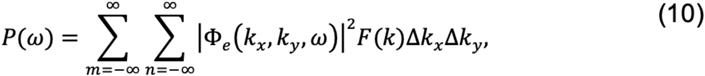

With

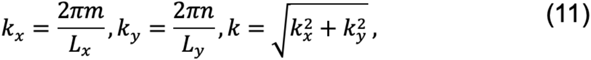

Where *L_x_* and *L_y_* are the spatial dimensions of the cortical sheet (*L_x_* × *L_y_*, with *L_x_* = *L_y_* = 0.5 *m*). The Gaussian filter function F(*k*) approximates the low-pass spatial filtering effects due to volume conduction, whose *k* dependence can be approximated by:

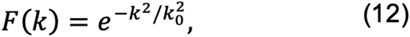

### Reduced model – Net circuit stability

The power spectral density is determined by five independent gain parameters (*G_ee_, G_ei_, G_ese_, G_esre_, G_srs_*). These five gain parameters can be further combined into three composite quantities that represent the effective gain within each of the model’s major feedback circuits, providing a low-dimensional physiological state space that characterises the net circuit stability ^30,72^:

Cortical-cortical gain:

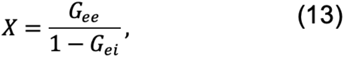

Corticothalamic loop gain:

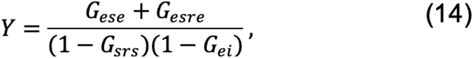

Intrathalamic loop gain:

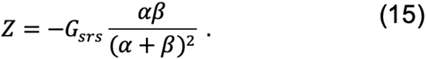

### EMG artifact model

To account for residual high-frequency electromyographic (EMG) contamination in the scalp EEG, an EMG component is added to the model-predicted spectrum before fitting:

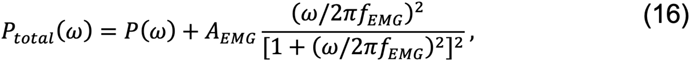

where *A_EMG_* is the EMG amplitude and *f_EMG_* is the characteristic frequency of the muscle noise contribution.

### Spectral and Parameter fitting

Model parameters were fitted to experimental EEG power spectra using Markov Chain Monte Carlo sampling with a Bayesian sampling-based Metropolis-Hastings (MH) algorithm ^32^. The quality of the model fit is determined by the weighted fractional difference between the model-generated power spectrum prediction *P*_j_(*x*) and the experimental power spectrum *P^exp^*:

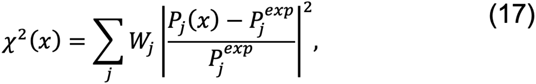

where *x* is the set of parameters to be estimated, *j* indexes frequency components from the fast Fourier transform of the experimental data, and the weighting *W*_j_ ∝ *f_j_*^-1^ acts as a scaling factor to provide equal weight to each frequency decade.

The MH algorithm estimates the parameter vector *x* that maximises the likelihood by performing a random walk through parameter space, evaluating the likelihood function *L* for each proposed parameter set,

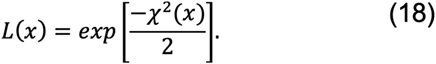

When modelling spectra for the frontal and parietal regions, no significant lateralisation effects were observed in the parameter estimates (p_FDR_ > 0.05). To simplify the analysis, we averaged parameters across both hemispheres. Examples of the model fits are shown in Figure S9.

## ACKNOWLEDGMENTS

C.N.R. is supported by an Australian Government Research Training Program Scholarship and a Living Stipend through the administration of the University of Queensland. C.N.R. is also supported by a QIMR-administered Living Stipend extension funded by L.J.H. This work was supported by the Australian NHMRC (LC: GN2001283 and GNT2027597, LJH: APP1194070).

## AUTHOR CONTRIBUTIONS

C.N.R. conceptualised the research goals, collected and pre-processed the data, formally analysed the data and wrote the first draft. L.J.H. and T.I. analysed the data and supervised the investigation. K.K.I. and J.A.R. supervised the neural field modelling. L.C. supervised the investigation and wrote the first draft. All authors interpreted the data, reviewed and edited the manuscript.

## DATA AVAILABILITY STATEMENT

- The EEG data will be made available to researchers following the execution of a data transfer agreement with QIMR Berghofer. Further information and requests for data should be directed to Luca Cocchi
- Analysis scripts to reproduce all figures are available on GitHub (released upon publication).
- Any additional information required to reanalyse the data reported in this paper is available from the lead contact upon request.

## SUPPLEMENTAL INFORMATION

Supplemental information can be found in Document S1. Figures S1–S9 and Table S1.

