## Supplementary_01 for "Complementary frontoparietal and corticothalamic contributions to relational reasoning"

---

**Supplementary Material**

---

**Contents:**

- **Supplementary Figure 1.** *Group-level power reliability across broadband and band-specific frequencies as a function of correct trials.*
- **Supplementary Figure 2.** *Right hemisphere frontal and parietal differences in regional cortical power as a function of increased relational complexity.*
- **Supplementary Figure 3.** *Left hemisphere frontal and parietal differences in induced and evoked activity as a function of relational complexity.*
- **Supplementary Figure 4.** *Right hemisphere frontal and parietal differences in induced and evoked activity as a function of relational complexity.*
- **Supplementary Figure 5.** *Aperiodic and periodic modelling of the spectral components of the post-stimulus relational integration window.*
- **Supplementary Figure 6.** *Mediation analysis modelling the brain-behaviour relationship between transitions in RC.*
- **Supplementary Figure 7.** *Model-derived engagement of the corticothalamic system during the encoding of relational complexity to the somatomotor cortex.*
- **Supplementary Figure 8.** *Cortical parcellations used to define regions of interest for EEG source localisation.*
- **Supplementary Figure 9.** *Example CTM spectral model fits to the empirical power spectrum during the relational integration window.*
- **Supplementary Table 1.** *Model parameters and constants.*
- **Supplementary References**

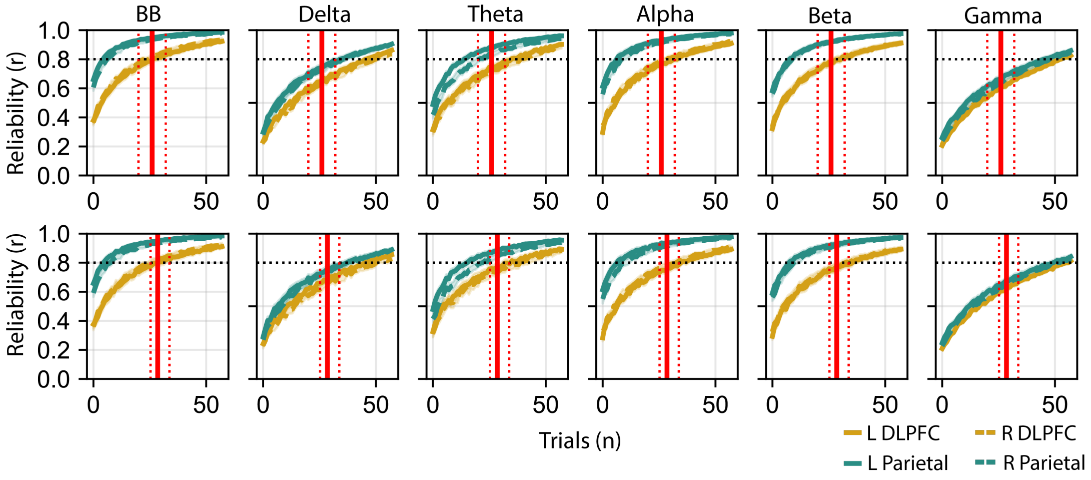


**Supplementary Figure 1. Group-level power reliability across broadband and band-specific frequencies as a function of correct trials.**

Visualisation limited to the first 60 sampled trials out of 192. Vertical red lines indicate the interquartile range (IQR) of minimum trials per condition: Top: All participants (n = 45, IQR: median = 26 [20 – 32] trials), Bottom: Participants included in final analyses after applying minimum trial criteria (n = 34, IQR: median = 28.5 [25.25 – 33.75] trials). Horizontal black line shows a correlation value of r=0.8, an arbitrary threshold indicating signal reliability [6]. Parietal regions (green) achieve stable power estimates with fewer trials than frontal regions (gold), with consistent patterns across hemispheres (solid vs. dotted lines). Using the study's minimum criterion of 20 trials yields reliable signals (r ≥ 0.8) in broadband, theta, alpha, and beta frequencies, while delta and gamma bands require cautious interpretation. From left to right: Broadband (BB: 2 – 45 Hz), Delta (2 – 4 Hz), Theta (4 – 8 Hz), Alpha (8 – 12 Hz), Beta (13 – 30 Hz), and low Gamma (30 – 45 Hz).

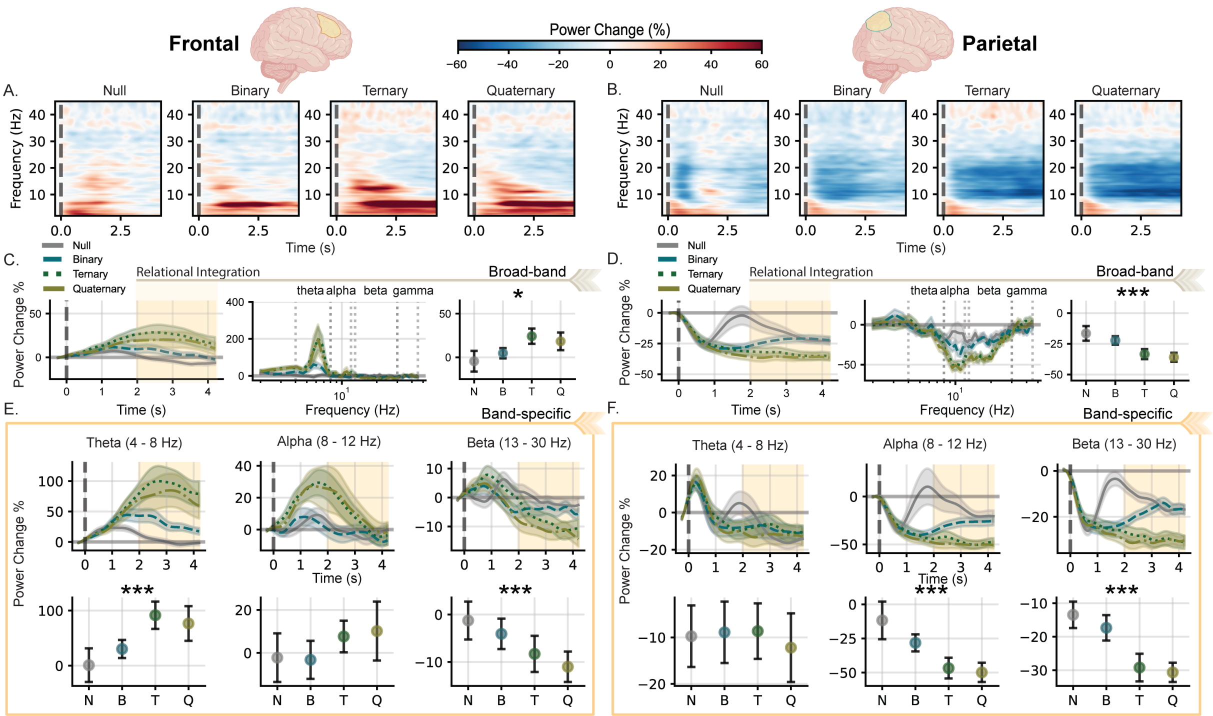


**Supplementary Figure 2. Right Hemisphere Frontal and Parietal differences in regional cortical power as a function of increased relational complexity.**

Group-level spectral power dynamics across frontal (left panels) and parietal (right panels) regions during relational processing (n = 34). (A, B) Time-frequency representations (2-45 Hz) showing percentage power change from pre-stimulus baseline (-0.3 to -0.05 s) across four complexity conditions (null, binary, ternary, quaternary). Warm colours indicate increases in power; cool colours indicate decreases in power relative to baseline. (C, D) Broadband power (2-45 Hz) time series across the puzzle stimulus, with shaded regions representing standard error of the mean. Baseline normalised power spectra (centre) extracted during the relational integration window (2-4.2 s, highlighted in beige[2]). Interval plots (right) display condition means with within-subject 95% confidence intervals. (E, F) Band-specific time-series for theta (4-8 Hz), alpha (8-12 Hz), and beta (13-30 Hz) frequencies, with mean activity during the relational integration window below. Asterisks indicate significant differences between conditions (*$p_{FDR} < 0.05$, ***$p_{FDR} < 0.001$).

Although our prior representational similarity analyses successfully decoded relational complexity at the single-trial level [2], that approach could not determine whether puzzles of the same complexity elicit a shared, time-locked neural response, rather than merely sharing relational structure. Figures S3 and S4 show the left and right hemispheres' induced and evoked responses to RC demands.

Induced left frontal broadband ($F(1.21, 39.80) = 4.94$, $p_{FDR} = 0.039$, $\eta_p^2 = 0.13$) and theta band ($F(1.27, 41.87) = 7.29$, $p_{FDR} = 0.014$, $\eta_p^2 = 0.181$) power increased, while beta band ($F(3, 99) = 10.22$, $p_{FDR} < 0.001$, $\eta_p^2 = 0.237$; Figures S3B and S3C *left*) power decreased, mirroring the total power results presented in Figure 2E. Likewise, decreases in induced power in the left parietal cortex occurred across the three frequency bands similar to the results presented in the total signal (theta: $F(2.08, 68.69) = 5.616$, $p_{FDR} = 0.003$, $\eta_p^2 = 0.145$; alpha: $F(1.36, 44.83) = 21.09$, $p_{FDR} < 0.001$, $\eta_p^2 = 0.390$; beta: $F(2.38, 78.66) = 56.00$, $p_{FDR} < 0.001$, $\eta_p^2 = 0.629$; Figures S3B and S3C *right*). Induced broadband and band-specific power changes in the right hemisphere (Figures S4B and S4C) also mirrored the total power results presented in Figure S2. On the other hand, across both hemispheres, we did not observe any power changes in the evoked signal during the relational integration window across complexity conditions in either frontal or parietal regions (Figures S3 and S4, left and right hemispheres, respectively).

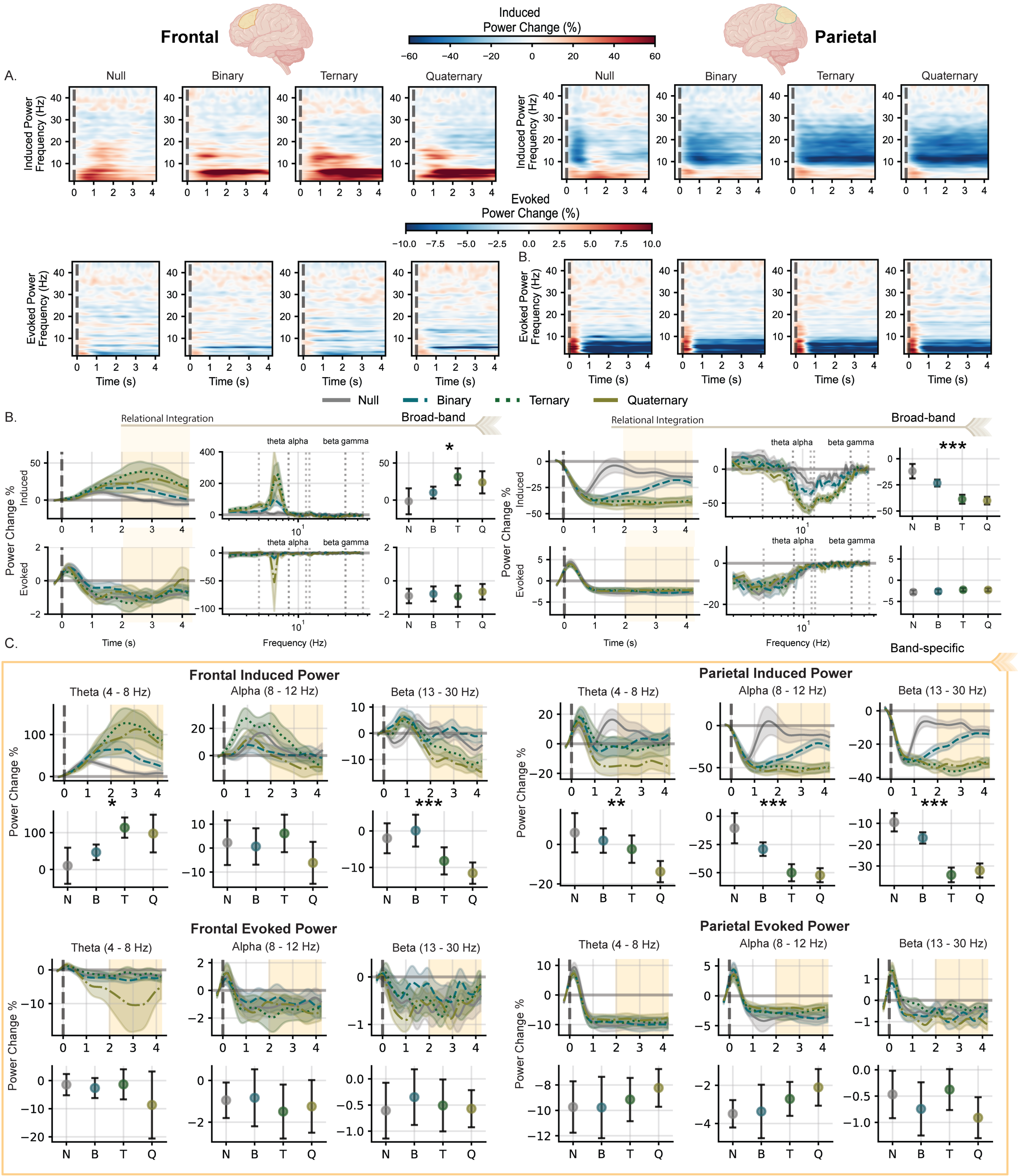


**Supplementary Figure 3. Left hemisphere frontal and parietal differences in induced and evoked activity as a function of relational complexity.**

(A) Time-frequency representations (2-45 Hz) showing percentage power change from pre-stimulus baseline (-0.3 to -0.05 s) across four complexity conditions (null, binary, ternary, quaternary). Warm colours indicate increases in EEG signal power; cool colours indicate decreases in power relative to baseline. (B) Broadband power (2-45 Hz) time series across the puzzle stimulus, with shaded regions representing standard error of the mean. Baseline normalised power spectra (centre) extracted during the relational integration window (2-4.2 s, highlighted in beige[2]). Interval plots (right) display condition means with within-subject 95% confidence intervals. (C) Band-specific time-series for theta (4-8 Hz), alpha (8-12 Hz), and beta (13-30 Hz) frequencies, with mean EEG signal power during the relational integration window below. Asterisks indicate significant differences between conditions (*$p_{FDR} < 0.05$, **$p_{FDR} < 0.01$, ***$p_{FDR} < 0.001$).

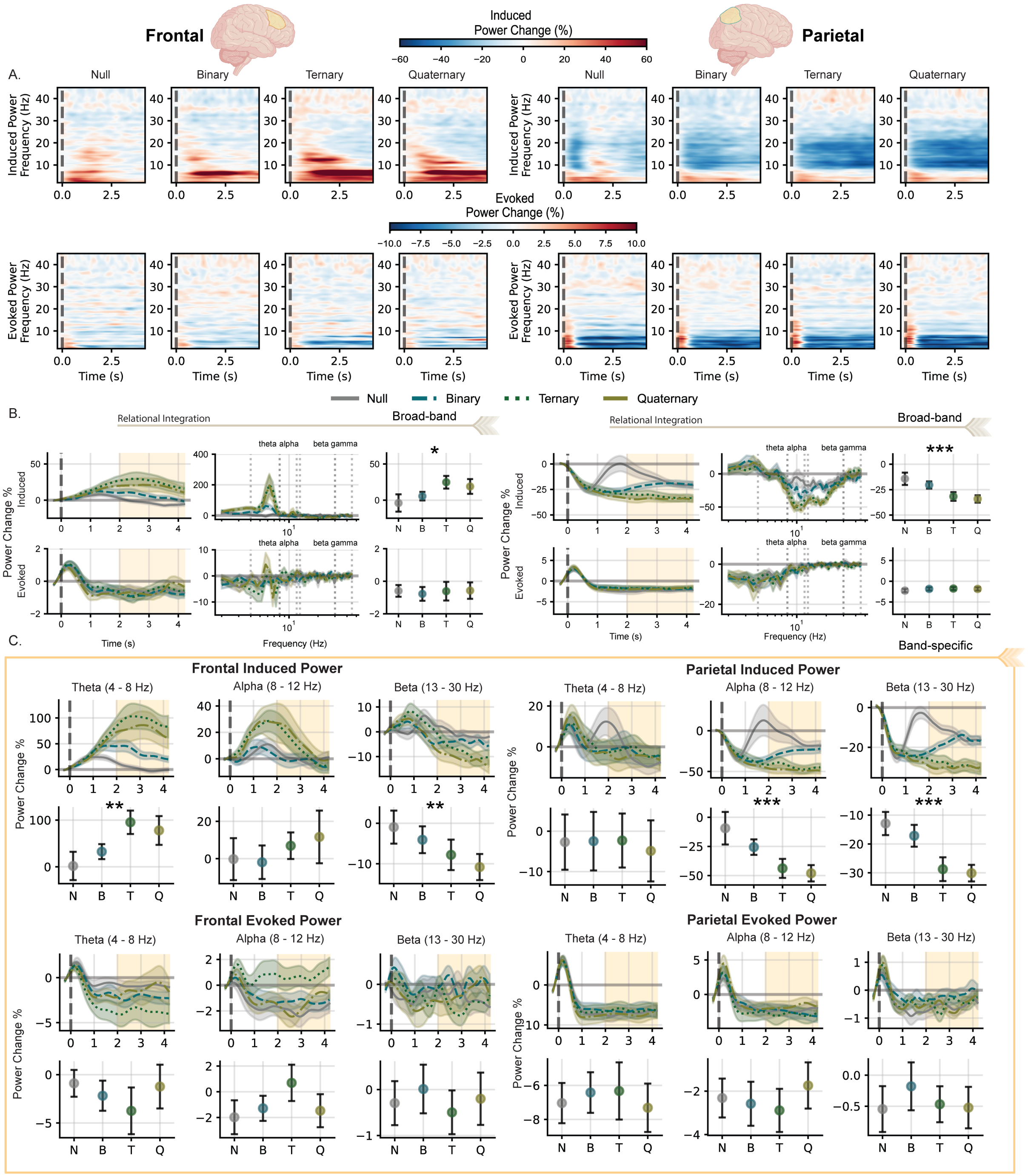


**Supplementary Figure 4. Right hemisphere frontal and parietal differences in induced and evoked activity as a function of relational complexity.**
(A) Time-frequency representations (2-45 Hz) showing percentage power change from pre-stimulus baseline (-0.3 to -0.05 s) across four complexity conditions (null, binary, ternary, quaternary). Warm colours indicate increases in EEG signal power; cool colours indicate decreases in power relative to baseline. (B) Broadband power (2-45 Hz) time series across the puzzle stimulus, with shaded regions representing standard error of the mean. Baseline normalised power spectra (centre) extracted during the relational integration window (2-4.2 s, highlighted in beige). Interval plots (right) display condition means with within-subject 95% confidence intervals. (C) Band-specific time-series for theta (4-8 Hz), alpha (8-12 Hz), and beta (13-30 Hz) frequencies, with mean EEG signal power during the relational integration window below. Asterisks indicate significant differences between conditions (*$p_{FDR} < 0.05$, **$p_{FDR} < 0.01$, ***$p_{FDR} < 0.001$).

In Figures S5A and S5C, in both hemispheres the frontal exponent and slope offset increased with relational load (left hemisphere: exponent: $F(3, 99) = 11.263$, $p_{FDR} < 0.001$, $\eta_p^2 = 0.254$; offset: $F(3, 99) = 10.193$, $p_{FDR} < 0.001$, $\eta_p^2 = 0.236$; right hemisphere: exponent: $F(3, 99) = 9.123$, $p_{FDR} < 0.001$, $\eta_p^2 = 0.217$; offset: $F(3, 99) = 9.181$, $p_{FDR} < 0.001$, $\eta_p^2 = 0.218$). In the left parietal cortex, aperiodic activity showed a complexity-dependent decrease in both the exponent and offset (Figure S5B: exponent: $F(3, 99) = 3.685$, $p_{FDR} = 0.018$, $\eta_p^2 = 0.100$; offset: $F(3, 99) = 8.44$, $p_{FDR} < 0.001$, $\eta_p^2 = 0.204$). The right parietal cortex showed a similar complexity-dependent decrease in the offset but no significant difference in the exponent (Figure S5D: exponent: $F(3, 99) = 1.560$, $p_{FDR} = 0.255$, $\eta_p^2 = 0.045$; offset: $F(3, 99) = 4.075$, $p_{FDR} = 0.015$, $\eta_p^2 = 0.110$). These results confirm that increasing complexity leads to distinct regional effects in aperiodic activity. Frontal areas exhibit an upward shift and steepening of the spectral slope, while parietal areas show the opposite pattern. Modelling the periodic, band-specific activity while accounting for aperiodic components revealed isolated periodic activity that persisted in both frontal and parietal areas. In the left and right frontal cortex, periodic spectra showed increased theta power with a concurrent decrease in beta activity (left hemisphere, Figure S5A theta: $F(3, 99) = 6.192$, $p_{FDR} < 0.001$, $\eta_p^2 = 0.158$; beta: $F(3, 99) = 12.414$, $p_{FDR} < 0.001$, $\eta_p^2 = 0.273$; right hemisphere, SI Appendix, Figure S5C theta: $F(3, 99) = 6.481$, $p_{FDR} < 0.001$, $\eta_p^2 = 0.164$; beta: $F(3, 99) = 13.317$, $p_{FDR} < 0.001$, $\eta_p^2 = 0.288$). The left and right parietal regions showed selective decreases in alpha and beta bands (left hemisphere, Figure S5B alpha: $F(3, 99) = 21.493$, $p_{FDR} < 0.001$, $\eta_p^2 = 0.394$; beta: $F(3, 99) = 62.301$, $p_{FDR} < 0.001$, $\eta_p^2 = 0.654$; right hemisphere, Figure S5D alpha: $F(3, 99) = 17.420$, $p_{FDR} < 0.001$, $\eta_p^2 = 0.345$; beta: $F(3, 99) = 23.463$, $p_{FDR} < 0.001$, $\eta_p^2 = 0.416$), with no significant modulation in theta power.

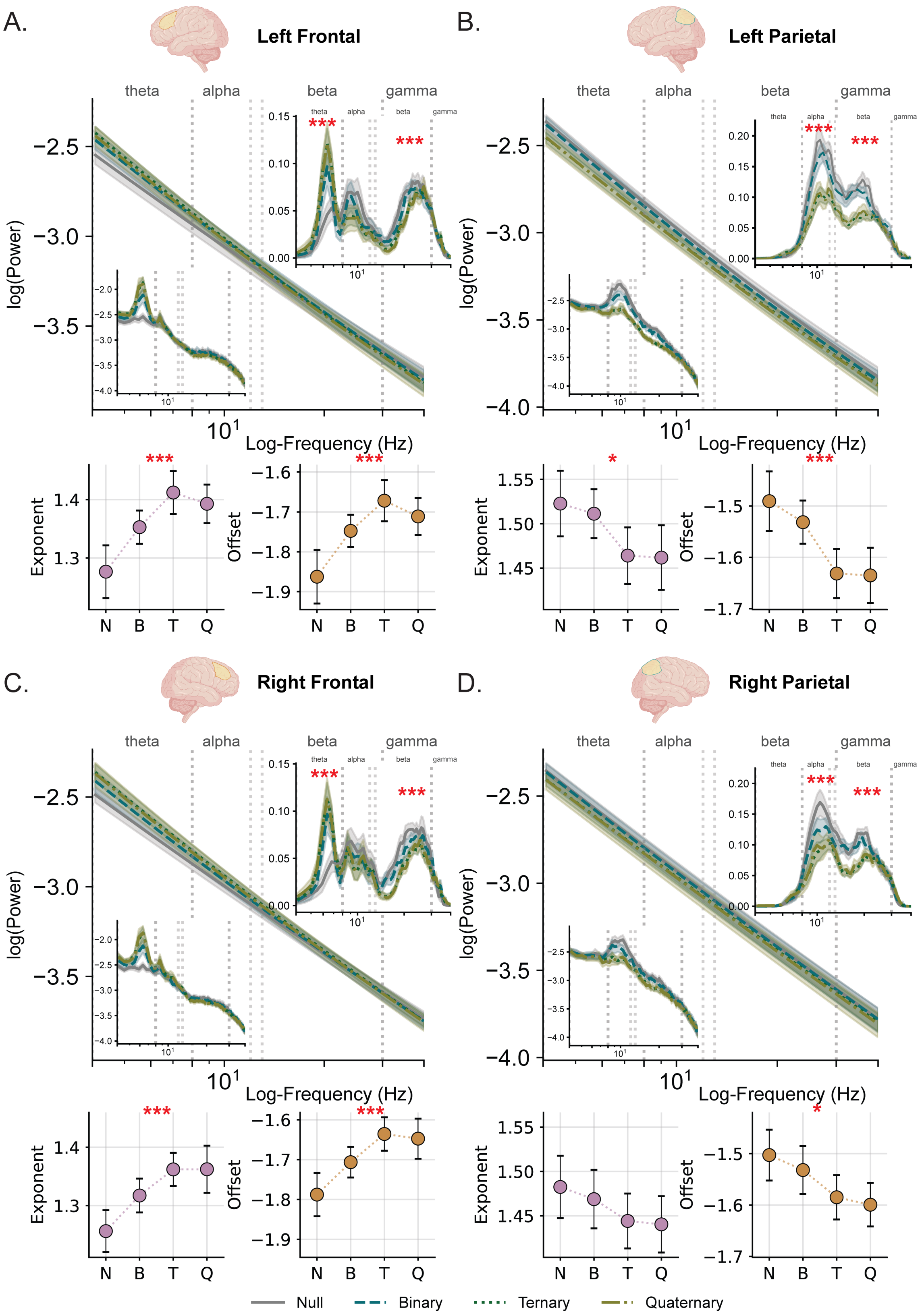


**Supplementary Figure 5. Aperiodic and Periodic Modelling of the spectral components of the post-stimulus relational integration window.**

Log-log plot of the group-averaged mean spectral aperiodic (main plot), raw spectra (embedded bottom left) and relative periodic power (embedded top right) components modelled between 4 - 40 Hz. Left hemisphere: frontal (A), parietal (B); right hemisphere: frontal (C), parietal (D). The shaded area depicts the standard error of the mean. Null (grey), Binary (blue), Ternary (dark green), and Quaternary (olive) conditions are shown for each frequency band. Red asterisks denote *p < 0.05, ***p < 0.001 for a one-way repeated measures ANOVA.

For the most extreme increase in RC (Model 1: Quaternary-Binary), scaling neural activity was specific to the theta band (Figure S6A). In this model, increases in frontal–parietal theta power were associated with increases in frontoparietal network connectivity (a = 0.155, CI [0.004, 0.305], p = 0.044).

For the more incremental increase in RC (Model 2: Quaternary-Ternary), beta-band activity revealed a dissociation between nodal power and network synchronisation in their relationship to behaviour (Figure S6B). Changes in beta network power did not predict beta-band synchronisation (a = -0.013, CI [-0.131, 0.106], p = 0.827). However, increases in network synchronisation were associated with longer response times (b = 8.073, CI [3.242, 12.093], p = 0.002) and reduced accuracy (b = -8.161, CI [-14.570, -1.754], p = 0.014). In contrast, increases in nodal beta power showed the opposite pattern, tending to correlate with faster response times (c′ = -1.331, CI [-2.877, 0.215], p = 0.089) and significantly improved accuracy (c′ = 2.863, CI [0.986, 4.740], p = 0.004).

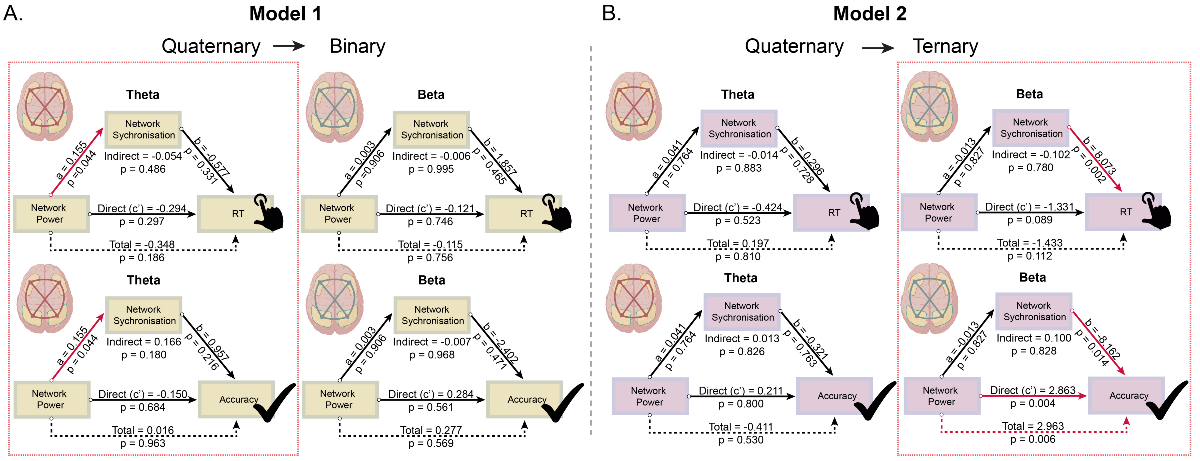


**Supplementary Figure 6. Mediation analysis modelling the FPN brain-behaviour relationship between RC demands.**
All Mediation models used to examine how changes in theta and beta band-specific network power and phase synchronisation covary with behaviour. (A) Model 1: the difference between low to high RC (Quaternary *minus* Binary). (B) Model 2: the difference between medium to high RC (Quaternary *minus* Ternary). Mediation analysis performed using a bias-corrected non-parametric bootstrap method (95%, CI 5000 iterations). Path coefficients and p-values are reported for each pathway (black arrows). Significant individual regression paths are indicated by red arrows (p < 0.05). The red box indicates the models reported in the main text Figure 3.

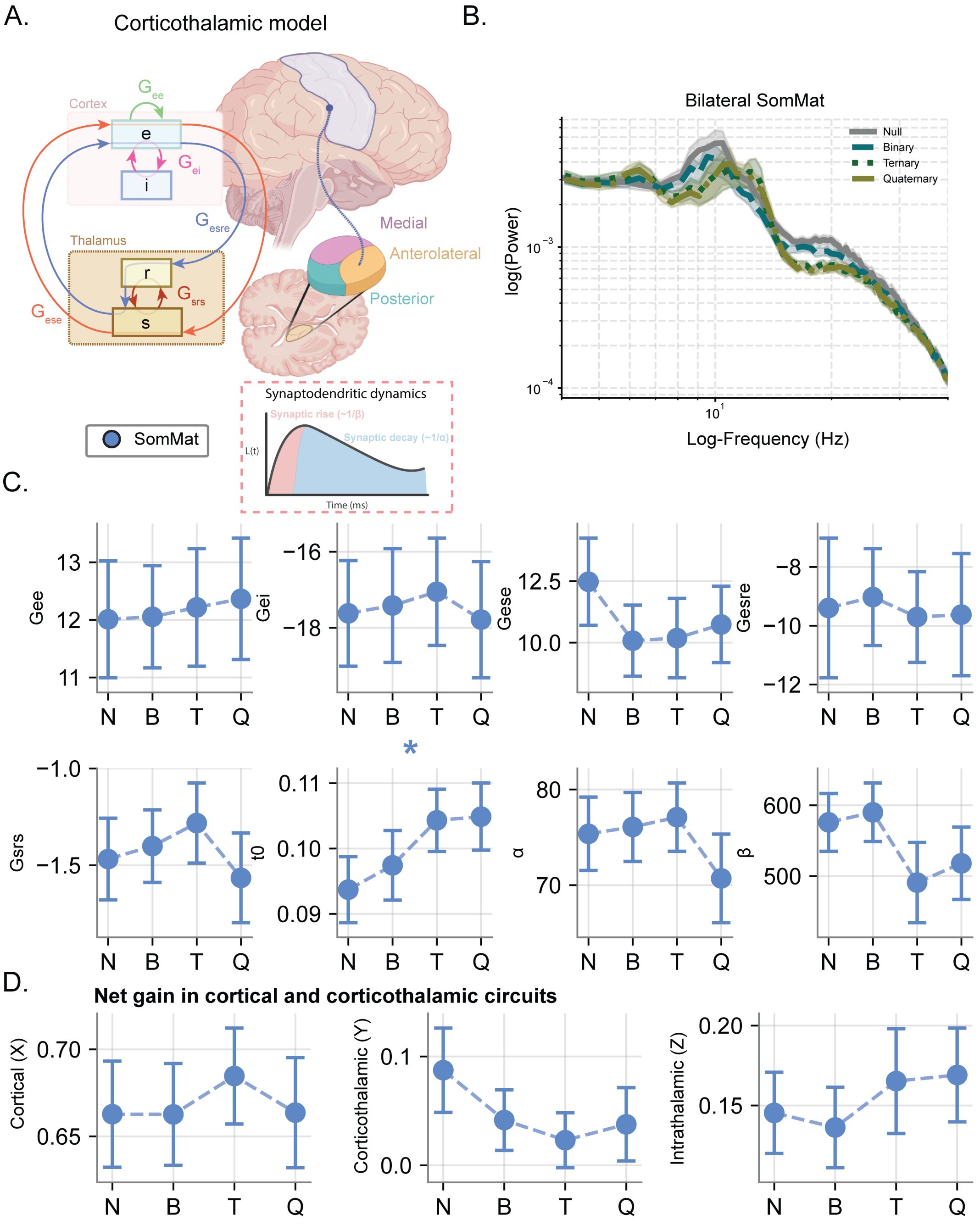


**Supplementary Figure 7. Model-derived engagement of the corticothalamic system during the encoding of relational complexity in the Somatomotor cortex.**

(A) Schematic of the corticothalamic model showing: cortical excitatory (e) and inhibitory (i) populations, inhibitory thalamic reticular (r) and excitatory thalamic relay neural populations (s), and their connections to the *Somatomotor* (blue) region. Synaptic decay $\alpha$ and rise $\beta$ describe the synaptodendritic dynamics. (B) Log-log plot of the power spectral density averaged across hemispheres across RC conditions. Error bars show mean and standard error of the mean. (C) Sub-circuit CTM parameters across increasing relational complexity levels (N = null, B = binary, T = ternary, Q = quaternary relations). (D) Net cortical, corticothalamic, and intrathalamic gain across complexity levels. Significant complexity effect *$p_{FDR} < 0.05$, Data show mean ± 95% within-subject confidence intervals.

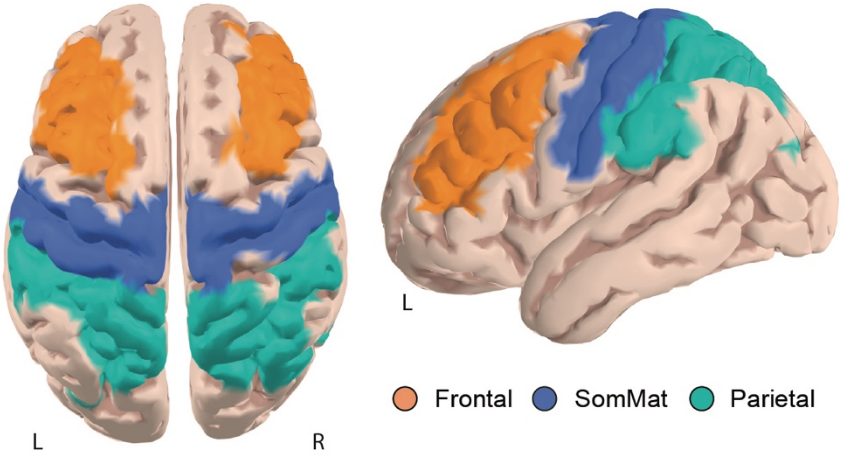


**Supplementary Figure 8. Cortical parcellations used to define regions of interest for EEG source localisation.**
Frontal, Parietal, and Somatomotor regions are projected onto the ICBM152 cortical surface template [7,12]. L = left, R = right.

Model fit quality was evaluated using the weighted chi-squared error ($\chi^2$; Equation 17), which quantifies the difference between the empirical and predicted power spectra. Following the criterion used in the original BrainTrak validation, fits with $\chi^2 < 4$ were deemed acceptable (23). Across all regions (Frontal, Parietal, Somatomotor), RC levels, and participants, the mean chi-squared was 0.66 (range 0.33–1.59).

To assess whether fit quality varied systematically with RC, a one-way within-subject repeated-measures ANOVA was conducted separately for each region. Fit quality did not differ significantly across conditions in frontal cortex ($F(2.15, 70.78) = 1.85$, $p = .162$) or parietal cortex ($F(2.26, 74.56) = 1.50$, $p = .228$). In contrast, fit quality in somatomotor cortex varied significantly with complexity, with relatively worse fits in ternary and quaternary ($F(2.36, 77.80) = 12.57$, $p < .001$). Because the somatomotor cortex served as a control region rather than a region of theoretical interest, this pattern does not affect the interpretation of the frontoparietal findings reported in the main analyses. Importantly, this reflects a relative reduction in the quality of fit. Overall, CTM provided a close characterisation of the empirical power spectra observed across the full range of task demands. Figure S9 shows representative examples of empirical and model-fitted power spectra for two participants (Subject 1, null condition, left parietal cortex; Subject 30, ternary condition, left frontal).

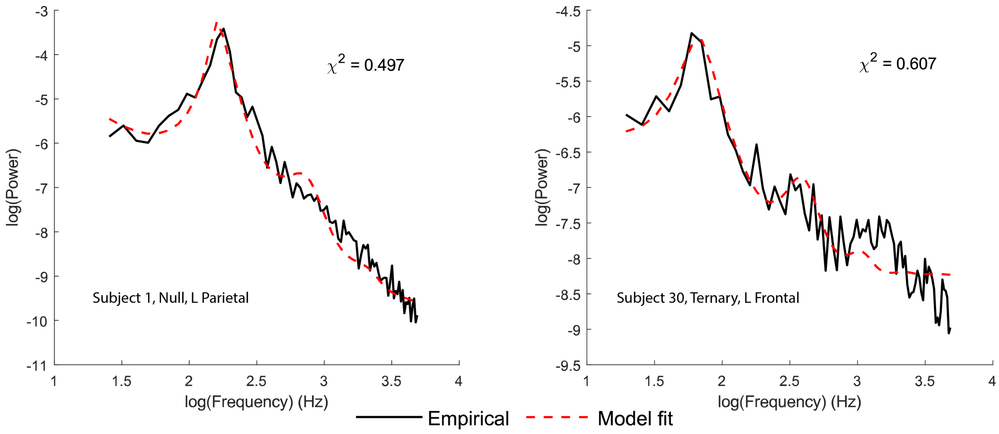


**Supplementary Figure 9. Example CTM spectral model fits to the empirical power spectrum during the relational integration window.**
Left, Subject 1, null condition, left parietal cortex ($\chi^2 = 0.497$). Right, Subject 30, ternary condition, left frontal ($\chi^2 = 0.607$). Black, empirical power spectrum; red dashed, model-fitted spectrum. Both examples illustrate representative model fits within the acceptable range ($\chi^2 < 4$;[23]).

**Supplementary Table 1. Model parameters and constants**

| Symbol | Description | Value | Range (Min, Max) | Initial Step | Unit |
|---|---|---|---|---|---|
| Model Parameters (to be fitted) | | | | | |
| $G_{ee}$ | Excitatory cortical gain | - | (0, 20) | 0.4 | - |
| $G_{ei}$ | Inhibitory cortical gain | - | (-40, 0) | 0.4 | - |
| $G_{ese}$ | Excitatory corticothalamic loop gain | - | (0, 40) | 1 | - |
| $G_{esre}$ | Inhibitory corticothalamic loop gain | - | (-40, 0) | 1 | - |
| $G_{srs}$ | Intrathalamic loop gain | - | (-5, 0) | 0.2 | - |
| $\alpha$ | Decay rate of cell body potential | - | (10, 100) | 5 | $s^{-1}$ |
| $\beta$ | Rise rate of cell body potential | - | (100, 800) | 40 | $s^{-1}$ |
| $t_0$ | Corticothalamic loop delay | - | (75, 140) | 5 | ms |
| $A_{EMG}$ | Normalization of EMG power | - | (0, 1) | 0.05 | - |
| Model Constants | | | | | |
| $\theta$ | Firing threshold | 12.9 | - | - | mV |
| $\sigma'$ | Threshold spread | 3.8 | - | - | mV |
| $Q_{max}$ | Maximum firing rate | 340 | - | - | $s^{-1}$ |
| $r_e$ | Excitatory axon range | 86 | - | - | mm |
| $\gamma_e$ | Cortical damping rate | 116 | - | - | $s^{-1}$ |
| $L_x, L_y$ | Cortical sheet length, width | 0.5 | - | - | m |
| $\Phi_n(k, \omega)$ | Input stimulus amplitude | $1\times10^{-5}$ | - | - | $s^{-1}$ |
| $\Phi_n^{(0)}$ | Steady state stimulus | 1 | - | - | $s^{-1}$ |
| $k_0$ | Volume conduction filter constant | 10 | - | - | $m^{-1}$ |
| $f_{EMG}$ | EMG center frequency | 40 | - | - | $s^{-1}$ |
